# Using BONCAT-FACS to probe the active soil microbial community during nitrous oxide production

**DOI:** 10.64898/2026.07.10.737762

**Authors:** Jonah Gray, Jennifer Harris, Jason Kaye, Estelle Couradeau

## Abstract

Nitrous oxide (N_2_O) is a potent greenhouse gas and is largely produced by incomplete denitrification. Although we know many of the microbial species that denitrify, we are still unable to reliably predict N_2_O production from soils. Recent work in microbial ecology has shown that when key microbes are considered as members of functional ensembles rather than isolated, the predictive power linking their activity to emergent properties increases dramatically. We hypothesized that the active microbial community during high N_2_O production would be taxonomically distinct from the inactive portion and increases in N₂O production rates would correlate more strongly with increased abundance across multiple active taxa than with dominance by a single active species. We conducted a microcosm experiment where agricultural soil was incubated in anaerobic vials for up to 15 hours while tracking N_2_O production. Using bioorthogonal non-canonical amino acid tagging paired with fluorescence-activated cell sorting and 16S rRNA amplicon sequencing (BONCAT-FACS-Seq), we probed the active subset of the microbial community throughout the incubation period. Analysis of 16S rRNA gene amplicons revealed that the active and inactive fractions contained distinct taxa, and the taxonomic composition of the active fraction shifted over time. We found that less than 1% of the microbial community was responsible for N_2_O flux rates as high as 3.84 µg N_2_O-N g dry soil-1 hr-1. The level of activity (median fluorescent intensity of active cells) correlated well with N_2_O production rates. The Ensemble Quotient Optimization for Microbiomes (mEQO) tool was used to identify an ensemble of eight organisms whose combined abundance best correlated with N₂O fluxes. Overall, our results reveal that N₂O fluxes are driven not by changes in a single taxon but by shifting ensembles of active microorganisms whose combined functional potential supports consistent emissions. This study applied a novel conceptual and methodological framework with a distinct focus on the active microbial community, rather than the entire community; if our observation that N_2_O flux rates are correlated with an ensemble of organisms is broadly confirmed, then framing denitrification as a community trait may increase predictability of this key process.

## 1 Introduction

Agriculture is the largest source of global anthropogenic nitrous oxide (N_2_O) (Tian et al. 2020), and accounts for at least 27% of total nitrous oxide (N_2_O) emissions (Fowler et al, 2015). Nitrous oxide is a potent greenhouse gas with 273 times the warming potential of carbon dioxide on a 100-year time scale (Calvin et al., 2023), and it contributes to ozone depletion (Ravishankara et al, 2009). Although N_2_O can be produced aerobically (Myrold, 2021) and as a byproduct of nitrification (Inatomi et al, 2019), most N_2_O is produced as an intermediate of canonical denitrification (Myrold, 2021). Crucially, it remains a challenge to predict N_2_O fluxes due to their high variability in space and time (Charteris et al., 2020). Despite knowing that many microorganisms have denitrification capabilities, it remains unclear which species are active and responsible for N_2_O fluxes while denitrification is occurring (Butterbach-Bahl et al., 2013, Shan et al., 2021).

Canonical denitrification is a form of anaerobic respiration where nitrate is sequentially reduced to nitrite, nitric oxide, nitrous oxide, and di-nitrogen, as opposed to chemodenitrification or fungal denitrification (Myrold, 2021). For denitrification to occur, soil must be oxygen depleted and have available nitrogen in the form of nitrate or nitrite and carbon substrates. Oxygen depletion arises from a combination of reduced diffusion from the atmosphere (e.g. in wet soils) or high consumption from cellular respiration of soil organisms. As denitrification progresses towards dinitrogen gas, the presence of residual O_2_ and other, higher oxidized forms of N inhibits the use of N_2_O for complete denitrification (Gaskell et al, 1981), and as soil pores open (e.g. due to drying), N_2_O is allowed to diffuse to the surface, creating ephemeral, high N_2_O fluxes. The priming effect, a change in soil organic matter (SOM) mineralization rates started by a carbon or nitrogen addition, can also influence N_2_O fluxes. Priming mechanisms can increase or decrease N_2_O emissions from SOM. Four current hypotheses outline how C and N additions can mediate N_2_O production. In each of these hypotheses, nutrient additions drive microbial communities to increase or decrease SOM mineralization which subsequently affects the amount of N_2_O derived from SOM-N. (Daly et al., 2024).

Abiotic conditions are a strong driver of denitrification (Myrold, 2021), but the composition of the microbial community can also affect rates and end products of denitrification (Philippot et al., 2013, Domeignoz-Horta et al., 2015). Denitrifiers are a taxonomically diverse group (Tiedje, 1994) and make up about 0.1-5% of the soil microbiota (Myrold, 2021). Complete denitrifiers have one form of all the genes required to completely reduce nitrate to di-nitrogen gas. These reduction genes are *narG* and *napA* for nitrate, *nirS* and *nirK* for nitrite, *norB* for nitric oxide and *nosZ* for nitrous oxide. Recently, greater *nosZ* microbial diversity has been discovered suggesting new roles of *nosZ* function besides energy production (Shan et al., 2021). This work highlights the importance of non-canonical denitrifiers, organisms with only some of the denitrification genes, role in denitrification and N_2_O emissions. A recent study from Pold et al., 2025 suggests that denitrification is a “community trait” as they found partial denitrification pathways to be more common than complete pathways across the globe. Since there is a diversity of organisms that can fully or partially denitrify, understanding the community response to conditions primed for N_2_O production will be important to developing and enacting N_2_O management practices.

When evaluating the community response to denitrification conditions, it is critical to focus on active microbes. Only 0.1-2% of total microbial biomass in soil may be active at any time, (Blagodatskaya and Kuzyakov, 2013), and that canonical denitrifiers only make up 0.1-5% of the microbial community (Myrold, 2021), we expect that up to 2% of the soil microorganisms could be active under optimal conditions for N_2_O production. Although the organisms with the potential or proven ability to take part in denitrification have been previously identified (Graf et al., 2014, Shan et al., 2021), we lack data on how the active microbial community, including organisms that may not directly participate in denitrification but perform other forms of anaerobic respiration, contributes to denitrification. Recent work has shown that when key microbes are considered as members of functional ensembles rather than isolated, the predictive power linking their activity to emergent properties of the microbiome can dramatically increase. This approach is particularly useful for denitrification, given that multiple microbial taxa are commonly required for denitrification (Pold et al., 2025, Shan et al., 2021, 2023, Wu et al., 2021). In this paper, we seek to fill this gap by examining the active microbial community during high N_2_O production in an agricultural soil, with this approach we reframe N_2_O production from a function of single microbes to a property of an active community, and we link microbial community changes with N_2_O production. Integrating methods, including physiological probing and phenotype-based separation, following the framework of Next Generation Physiology (Hatzenpichler et al., 2020), will allow us to take the first step in linking microbial functions to soil emergent properties. This study will serve as a proof-of-principle to test this novel conceptual and methodological framework.

Our first hypothesis is that the active microbial community during high N_2_O production is taxonomically distinct from the inactive portion. Our second hypothesis is that increases in N_2_O production rates will correlate better with changes in the members of the active microbial community rather than changes in one active species (e.g. a complete denitrifier) dominating this metabolic pathway, indicating that the total flux of N_2_O is driven by a community response.

To test these hypotheses, we utilized biorthogonal non-canonical amino acid tagging (BONCAT) paired with fluorescent activated flow cytometry (FACS) to distinguish the active soil microbial community. BONCAT-FACS has been proven to be a powerful tool for analyzing the activity in soil microbial communities (Couradeau et al., 2019, Harris et al., 2025, Trexler et al., 2023). Using BONCAT-FACS, we analyzed how the active soil microbial community responded to optimal conditions for N_2_O production during a 15-hour lab incubation. We tracked the percentages of active and inactive cells, the cell abundances per gram of soil (Supplementary Figure 4), the median fluorescence intensity of cells, and physically sorted the active from inactive cells. We created a 16S rRNA gene library to compare the active community composition over time. By coupling our N_2_O data with the Ensemble Quotient Optimization for Microbiomes (mEQO) tool, we were able to identify an ensemble of organisms whose abundances correlate best with N_2_O production rates (Shan et al., 2023). We also employed the **P**hylogenetic **I**nvestigation of **C**ommunities by **R**econstruction of **U**nobserved **St**ates (PICRUSt2) software to predict functional gene abundances based on 16S amplicon sequences (Douglas et al., 2020) for the entire microbial community and the ensemble predicted by the mEQO tool. These data provide rare insights into links between microbial community change and N_2_O production.

## 2 Methods

### 2.1 Soil sampling and preparation

The soil used in this experiment was collected in October 2023 from an alfalfa field at The Russell E. Larson Agricultural Research Center in State College, Pennsylvania. We collected 5 kg of soil from the top 5 cm of soil. Each kg of soil was collected from field edges of two different alfalfa fields separated by a grass alleyway. We selected this specific soil because it was in an alfalfa production system for at least 2 years, so we expected it to have a relatively high fraction of nitrogen cycling microorganisms compared to soil from other crop production systems. Additionally, this soil is from the Hagerstown soil series, which is common in prime agricultural land of the Mid-Atlantic region of the United States (Soil Survey Staff, 2026).

All the collected soil was mixed, bagged, and refrigerated at 4 °C until the experiment in May 2024. Two subsamples of the remaining 5kg were stored at 4 °C or air dried for lab analysis and preservation. Lab analysis was done by the Pennsylvania State University Agricultural Analytical Services Laboratory in November 2025 to determine pH, organic matter, total carbon and nitrogen content, and texture. It has a pH of 6.7, an organic matter (OM) percentage of 2.97%, a total nitrogen percentage of 0.21%, and a total carbon percentage of 2.05% (C:N ratio ∼ 10). The soil is 28.1% sand, 44.0% silt and 27.9% clay, and is classified as a clay loam. Initial gravimetric water content was determined by weighing 10 g of fresh soil into a pre-weighed tin. Soil was dried in the oven at 105 °C for 2 days then reweighed. Before the experiment, the soil was homogenized using a number 5 sieve (4mm) to maintain small aggregates and to exclude aggregates larger than 4 mm that would not fit into exetainer vials.

The water-holding capacity was determined in April 2024, prior to the initiation of the incubation experiment. To determine average water holding capacity, 10 g of oven dried soil was added to funnels with pre-wetted filters and replicated 10 times. 50 ml of deionized water was poured gently over the soil and allowed to drain over 6 hours. Weight of wet soil was determined by subtracting the funnel and wet filter weight. 100% water holding capacity was determined using this equation:

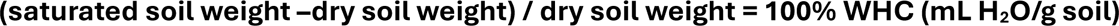

### 2.2 HPG solution preparation

Homopropargylglycine (HPG) is a non-canonical amino acid and methionine anolog used in BONCAT experiments. Thus, translationally active cells will utilize HPG instead of methionine to synthesize proteins. We used a Copper-catalyzed azide-alkyne cycloaddition (CuAAC) click chemistry reaction to “click” a fluorescent dye onto HPG-proteins as part of the BONCAT process. Additional preparation methods are in the supplemental materials.

### 2.3 Microcosm experiment

Five groups of 12 microcosm vials were incubated for varying times ranging from 3-15 hours. In each group, five vials were for BONCAT-FACS analysis, one for a water control for BONCAT-FACS, one contained autoclaved soil to serve as a background soil fluorescence control for BONCAT-FACS, and the remaining 5 for extractable nitrate and ammonium analysis. We incubated the microcosms for 3, 6, 9, 12, and 15 hours (supplementary figure 1A).

Prior to the incubation, two grams of fresh or autoclaved soil were added to each autoclaved 12.5 ml exetainer vial and stored in the refrigerator overnight. Samples were removed from the fridge and allowed to reach room temperature before a solution of glucose and potassium nitrate with a C:N mass ratio of 14 was added to achieve sixty percent water holding capacity. Vials were sealed with caps that included a septum to allow gas sampling. Then, using N_2_ gas, vials were purged of oxygen for 45 seconds to create anaerobic conditions. This was determined to be sufficient for maximizing N_2_O production over a 15-hour incubation.

Immediately following headspace purging, vials were placed into a dark incubator at 25 °C. For each incubation group, HPG was added via solution for the last 3 hours of incubation. Concentrated HPG solution was added to reach 95% water holding capacity. To limit O_2_ contamination, gas was sampled from an O_2_ free bottle and injected into the bottle containing O_2_-free HPG to keep HPG bottles pressurized. Then, HPG bottles were inverted to remove the necessary amount of HPG solution. Once HPG solution was removed from the bottle, it was added to the soil through the septum on the vial ensuring vials would remain anaerobic. HPG was sprinkled over the soil for even distribution.

Following each group’s incubation period, 7 of the 12 vials (5 BONCAT samples + 1 water control + 1 autoclaved soil control) had cells extracted. Cells were extracted in 1X phosphate buffer solution (PBS) with 0.02% Tween20. After vortexing for 5 minutes, vials were centrifuged at 500 RCF for 5 minutes at 4 °C. 2880 µl of supernatant was aliquoted evenly into 4 1.5 mL autoclaved Eppendorf tubes each prepared with 480 µL of 50% glycerol. After adding the supernatant, the concentration of glycerol was 20% so cell extracts could be frozen without cell lysis. Cell extracts were vortexed and frozen at -20 °C for later use. The remaining 5 vials had 2 M KCl added before storage at 4 °C for later use in nitrate and ammonium extractions.

### 2.4 Nitrous oxide, nitrate, and ammonium measurements

Nitrous oxide was measured using a closed loop apparatus in conjunction with a LI-7820 N_2_O trace gas analyzer (LI-COR Biosciences, 2023). Data were converted from ppb to µg N_2_O-N using the volume of the closed loop, the room temperature, atmospheric pressure and the ideal gas law. Since incubated vials had to be destructively harvested for cell extraction and BONCAT-FACS analysis, measurements of N_2_O occurred twice during each vial’s lifespan. Measurements were taken at 1 hour and ∼3 hours after HPG was added. Using a syringe, 0.25 ml of gas was removed for measurement. Flux rates were calculated over each 2-hour interval using the N_2_O-N measurements from each vial. Nitrate and ammonium concentrations were determined using a spectrophotometer and the Vanadium (III) Chloride method (Doane and Horwath, 2003) and the modified Indophenol Blue Method using salicylate (Utomo et al., 2023). Additional details can be found in the supplementary materials.

### 2.5 BONCAT-FACS: Cell sorting and cell counts

To prepare cell extracts for sorting for analysis on the Bigfoot Spectral Cell Sorter, cell extracts were thawed and pelleted (Supplementary Figure 1B). Most extracts still had low levels of soil particles and were first re-centrifuged at 500 RCF to remove excess soil particles. Then, the supernatant was transferred to a fresh, autoclaved Eppendorf tube. Cells were pelleted by centrifuging at 14000 RCF for 5 minutes. The supernatant was discarded, and cells were resuspended in 300 µL of sterile 1X PBS.

Following resuspension, cells underwent the “click” reaction with a solution containing the fluorescent dye, AF647-picolyl azide (excitation: 650 nm; emission: 670 nm), which indicates a BONCAT positive result in this study. This dye was selected as the BONCAT positive stain since the emission spectrum does not overlap with the emission spectrum of the counter stain, SYBR green (excitation: 498 nm; emission: 522 nm), a general DNA stain. Details on the preparation of the reaction mixture and samples can be found in the supplementary materials.

All cell sorting to determine inactive and active microbial populations took place between 7-17-2024 and 8-19-2024. Cell sorting used a nested gating strategy (Supplementary Figure 1C). SYBR stained samples were used to gate the “viable cell fraction” and exclude any remaining soil particles. The nested BONCAT gates for inactive and active cells used the AF647-picolyl azide dye. We gated against water controls (soils incubated with water instead of HPG) to determine the active fraction of cells and control for the false positive discovery rate (< 0.05%). Further cell sorting details can be found in the supplementary materials.

We correlated N_2_O fluxes with both the percentage of active organisms and the log of the median fluorescence. The log of the median fluorescence value is collected for each sample by the flow cytometer and indicates the median fluorescence of the fluorescing cells. Leizeaga et al., 2017 found that this value correlates with protein synthesis rates better than the percentage of cells that are active. To reduce batch effects in our calculations, we adjusted gates against our controls for each day a sort occurred. We calculated median fluorescent intensity by subtracting the median fluorescence collected from a water control with both fluorescent dyes present. The water control, in which cells incorporated no HPG, served as a control to estimate off-target fluorescent tagging in other samples run in the same batch and to limit our false-positive rate to less than 0.05%.

### 2.6 PCR and 16S rRNA gene amplicon data analysis

We used the MicroGEM prep kit, following the manufacturer’s recommendation, to extract and amplify DNA from as few as 20,000 cells. Low-yielding samples were mainly from the active fractions, and each sample’s active technical replicates needed to be combined to achieve higher cell counts of >30,000 cells. Therefore, the amplification of the active fraction was obtained by combining multiple aliquots collected from each replicate. We prepared our PCR mixture using the Invitrogen Platinum Master Mix, a GC enhancer, sterile water, and primers 515F and 806R (Apprill, et al., 2016, Caporaso et al., 2011, Caporaso et al., 2012, Parada et al., 2016) (forward primer: 5′- GTGYCAGCMGCCGCGGTAA-3′; reverse primer: 5′- GGACTACNVGGGTWTCTAAT-3′) to amplify the V4 region of the 16S rRNA gene sequence.

Once the 16S rRNA gene sequences were amplified, we sequenced them on an Illumina MiSeq sequencer with a 250 x 250 base pair paired-end run that returned about 65000 pairs per sample at the Huck Institute for the Life Science Genomics Core Facility. Data analysis was conducted in R using a DADA2 pipeline (Callahan et al., 2016). Additional details on the PCR preparation and 16S amplicon analysis can be found in the supplementary materials.

### 2.7 mEQO analysis

We utilized the mEQO (Ensemble Quotient Optimization for microbiome) R package that is designed to identify a group of species whose combined abundances correlate with a trait of interest (Shan et al, 2023). Since our trait of interest, nitrous oxide flux rate, is continuous, we could use either the Genetic Algorithm (GA) or Boolean Least Squares (BLS) algorithm ensemble quotient optimization. We opted for the Boolean Least Square algorithm due to its ease of use, speed, and lack of a priori knowledge about which ASVs should be included in the final ensemble. To use this tool efficiently, we needed to reduce the complexity of our ASV dataset. We achieved this by filtering out rare and uncommon ASVs across replicates and time points. We limited our dataset to only active ASVs that were present in the majority (60%) of all samples resulting in 537 ASVs being used in the analysis.

Using the BLS algorithm, we ran models that identified the best fit ensembles for 1-10 ASVs. We stopped at 10 ASVs to maintain accuracy and efficiency in fitting the model. Based on these 10 different models’ outcomes, we compared AIC (Akaike Information Criterion) values for each and selected the simplest model (lowest number of ASVs) with the lowest AIC value. This model included 8 ASVs. The 8 ASVs retrieved were further inspected using BLAST on their reference sequence. None of the sequences were associated with organisms that had full genome sequences, therefore further studies will be needed to identify their functional potential.

### 2.8 PICRUSt2 analysis

We employed the **P**hylogenetic **I**nvestigation of **C**ommunities by **R**econstruction of **U**nobserved **St**ates (PICRUSt2) software to predict functional gene abundances based off of 16S amplicon sequences (Douglas et al., 2020). We followed the pipeline outlined by creators with no changes, using our ASV table of abundances and 16S rRNA gene amplicon sequences as inputs. We opted to use the predicted samples gene family profiles output rather than the pathway abundances so we could identify abundance changes in specific nitrogen cycling genes rather than changes in the entire nitrogen metabolism pathway. We used these abundances to calculate the relative abundance of each gene (represented as a Kegg Orthology code) (Kanehisa and Goto, 2000) in each sample. Then, we used the Kegg Orthology codes to search for nitrogen cycling genes in the PICRUSt2 predicted genes. We searched for nitrogen fixation (*nifH*), nitrification (*amoA, amoB, amoC, hao*), dissimilatory nitrate reduction to ammonia (DNRA, *nrfA*), and denitrification (*napA, narG, nirK, nirS, norB, nosZ*) genes. All genes were predicted in our samples except for *amoA*. Genes *amoB and amoC* were predicted in our samples in very low relative abundance. Since these genes tend to be found in the *amoCAB* operon (Norton et al., 2001; Junier et al., 2009), we can expect the *amoA* gene should also be predicted in these samples. Analysis and interpretation will focus on *amoB* and *amoC.* Using the relative abundance of nitrogen-cycling genes in each sample, we used linear models to test for significant differences in predicted gene content between inactive and active fractions over time. Each model followed the same form: *log(predicted gene relative abundance + x) ∼ activity fraction * time*, where x is a small number added prior to log transformation to accommodate zero values and was defined as half of the smallest nonzero relative abundance value in the dataset. Activity fraction was treated as a categorical variable representing predicted genes from active or inactive samples, and the time point represented the incubation harvest time for each sample. Relative abundance values were log-transformed to improve normality and homogeneity of variance of model residuals.

## Results

### 3.1 Nitrogen cycling

Over the 15-hour incubation period, the N_2_O-N production rate (µg N_2_O-N g dry soil^-1^ hr^-1^) increased linearly (*N_2_O-N production rate = 0.19 + 0.11(hour), R^2^ = 0.94, p<0.001*;Figure 1) The maximum production rate, 3.84 µg N_2_O-N g dry soil^-1^ hr^-1^ was achieved between hours 13 - 15 of the incubation. Nitrate concentration decreased linearly during the incubation. (*mg NO_3_^-^-N/kg soil = -10.26 (hour) +189.99; R^2^=0.94, p <0.001*; Figure 1). On average, there was a 4.4-fold decrease in nitrate concentration from hour 3 to hour 15. Ammonium concentrations increased following a linear trend with incubation time (*mg NH_4_^+^ -N / kg soil= 0.16(hour) + 3.38; R^2^ = 0.58, p < 0.001;* Figure 1). While this represents a 50% increase in ammonium concentrations from the concentration at hour 3, ammonium concentrations were very low throughout the incubation, relative to nitrate.

**Figure 1:**
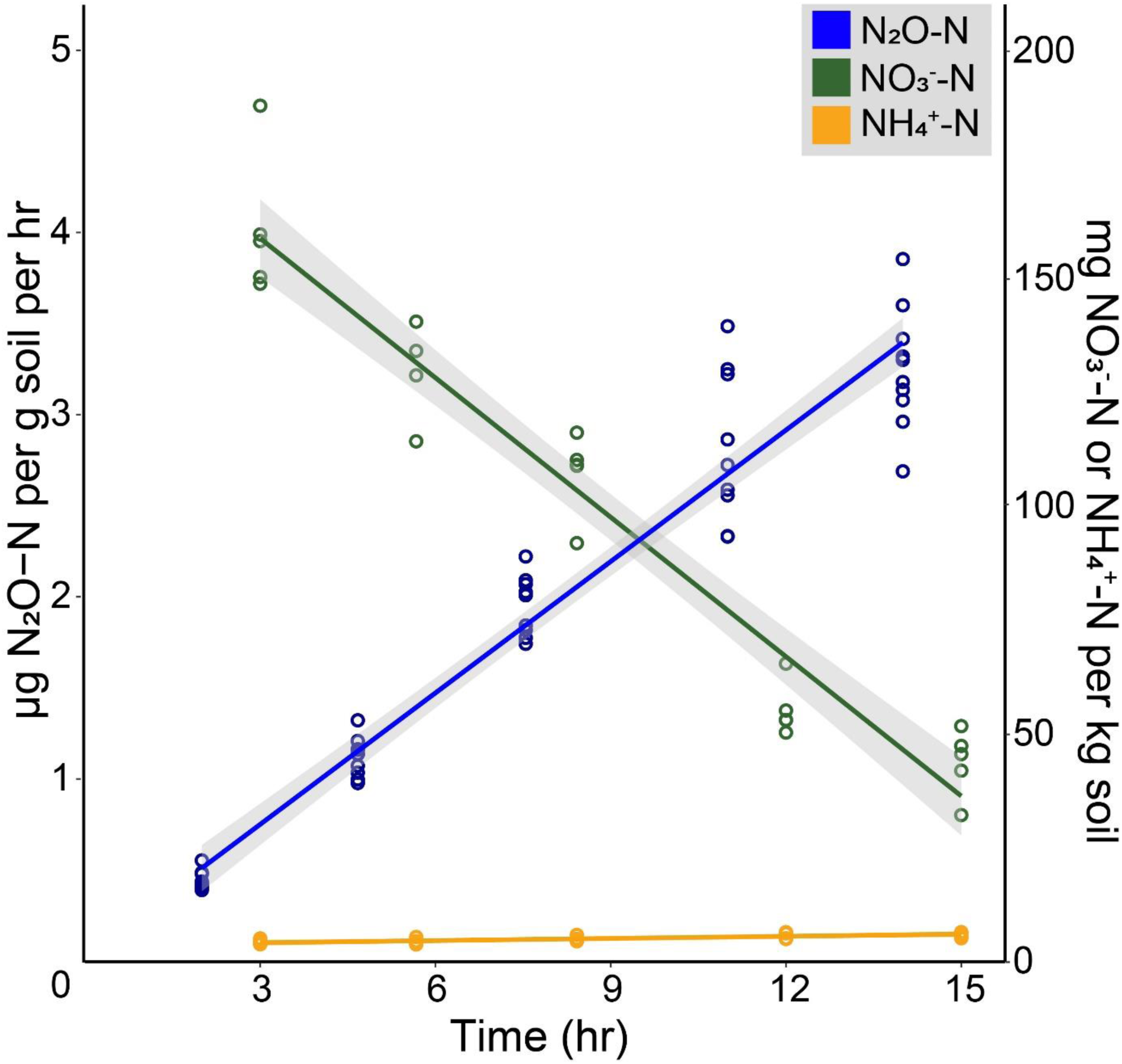
Changes in concentrations of key nitrogen species during the incubation. Nitrous oxide flux rates (blue, µg N_2_O-N g dry soil^-1^ hr^-1^) increase linearly (*p<0.001*) while nitrate (green, mg NO_3_^-^-N kg^-1^) concentrations decrease linearly (*p < 0.001*). Ammonium concentrations (orange, mg NH_4_^=^-N kg^-1^) increase linearly (*p < 0.001*).

### 3.2 BONCAT-FACS results

While N_2_O-N production rates increased, the percentage of cells that were active also increased, but never exceeded 1% (Figure 2A). The highest percentage of active cells was 0.79% at the fifth and final sampling time. However, the estimated mean active cells at this time point was 0.31%. Across the time points, we used a one-way ANOVA to test for overall differences. A significant difference was detected (F = 5.97, *p < 0.01,* df = 4), so we performed Tukey post-hoc comparisons to identify that the fifth time point is significantly different from the first three time points (*5-1, p < 0.01; 5-2, p < 0.01; 5-3, p < 0.01*; Figure 2A).

**Figure 2:**
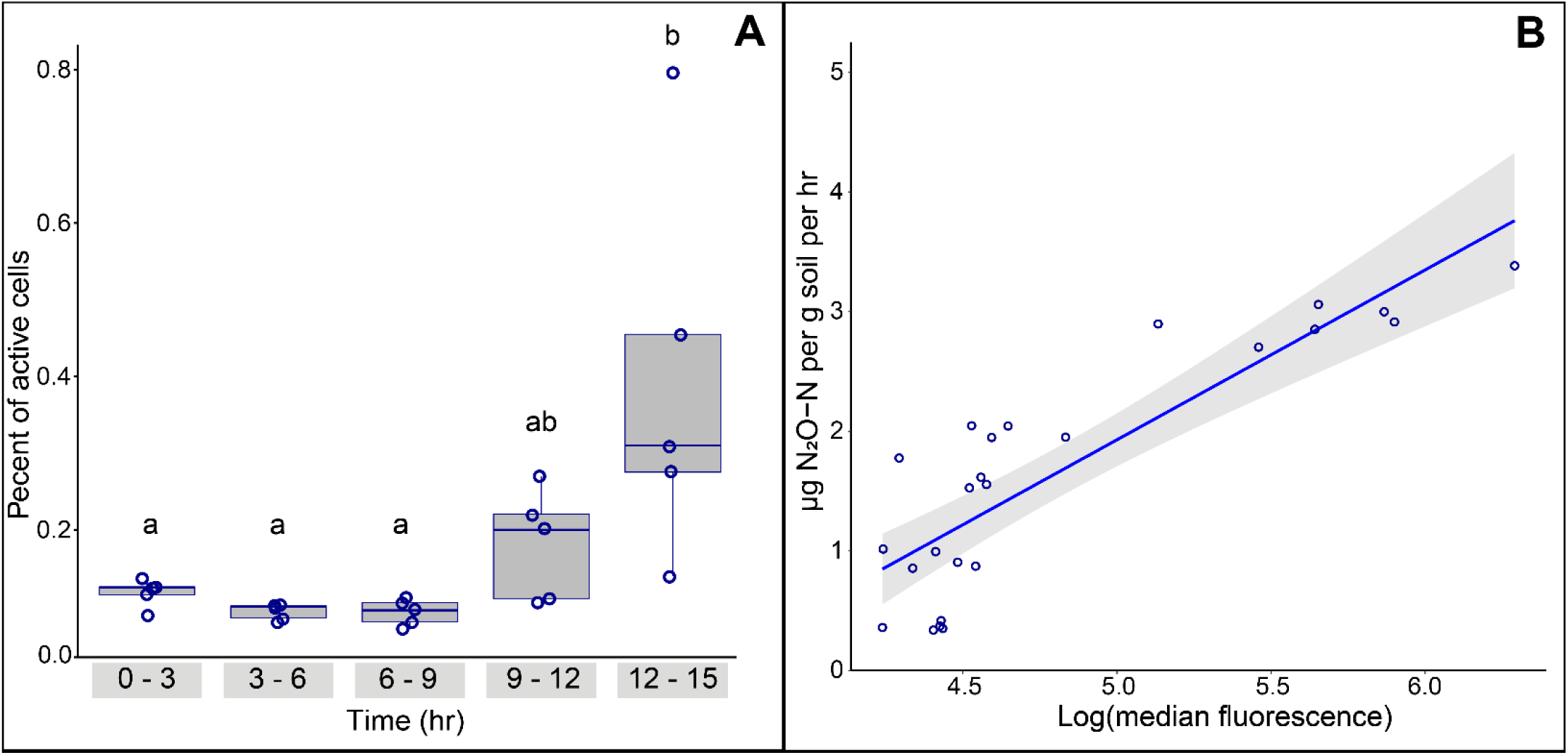
(A) Percentage of cells from the harvested soil samples that were active during each 3-hour time period. All time points had less than 1% activity. (B) Log(median fluorescence) had a linear relationship (*R^2^=0.74, p < 0.001*) with nitrous oxide flux rates (µg N_2_O-N g dry soil^-1^hr^-1^).

During the incubation, total viable cell count, estimated using the flow cytometer at a specific flow rate for a set time, did not change significantly over time (Supplementary Figure 4*, R^2^ = 0.03, p = 0.38*). This indicates that there was no net cell growth or death during the 15-hour incubation.

Although the percentage of cells that were active correlated with N_2_O production rates (*N_2_O-N production rate = 1.00 + 3.79(percent active); R^2^=0.33, p < 0.01*), the log of the median fluorescence correlation was stronger (Figure 2B*, N_2_O-N production rate = -5.17 + 1.42(log(median fluorescence)); R^2^=0.74, p < 0.001*). The log of the median fluorescence can be interpreted as a relative measure of how many proteins are being translated in a cell (Leizeaga et al., 2017).

### 3.3 Microbial community composition

Across samples, the observed number of ASVs ranged from 610 to 2265. We used a one-way ANOVA to test for differences in observed ASVs across (active, inactive, viable cells, and bulk soil). A significant difference was detected (Supplementary Figure 2, F = 17.98, *p < 0.001,* df = 3), so we employed Tukey post-hoc comparisons to determine that the number of observed ASVs from all viable cells and from the inactive fraction were both had significantly more observed ASVs than the active fraction (Supplementary Figure 2, *p < 0.001, p < 0.001*). The number of ASVs in the bulk soil was not significantly different from any of the other sample types. Following the same method, we detected a significant difference in Pielou’s evenness across samples (Supplementary Figure 3, F = 12.43, *p < 0.001,* df = 3) and found that evenness in the active fraction was significantly lower than all viable cells and the inactive fraction (Supplementary Figure 3, *p < 0.001, p < 0.001*) and that the bulk soil was not significantly different in evenness than any other fraction. We analyzed evenness over time in the active fraction and a one-way ANOVA showed a significant difference over time (Supplementary Figure 3, *p < 0.001)*. We investigated further using a Tukey post-hoc test that showed that the last two time points were significantly different than the first three (Supplementary Figure 3, *4-1, p < 0.001; 4-2, p < 0.001; 4-3, p < 0.001; 5-1, p < 0.001; 5-2, p < 0.001; 5-3, p < 0.001*), and that the first three time points were not significantly different from each other nor were the last two time points significantly different from each other.

The ordination of the Bray Curtis beta diversity dissimilarity metric using a PCoA showed that the active (red), bulk soil (blue), and inactive communities (green) were distinct (Figure 3), and axis 1 and axis 2 explain 16% and 5.7% of the total variation, respectively. Overall, this is relatively low amount of variation explained by both axes which may limit interpretation. A PERMANOVA analysis revealed that the activity status, time point, and the interaction between the two were all significant predictors of the variation in community composition (F = 7.07, *p < 0.001,* df = 3; F = 3.31, *p < 0.001,* df = 1; F = 2.40, *p < 0.001*, df = 2). This analysis showed that 21% of the variation in community composition can be explained by fraction (viable, bulk, active or inactive).

**Figure 3:**
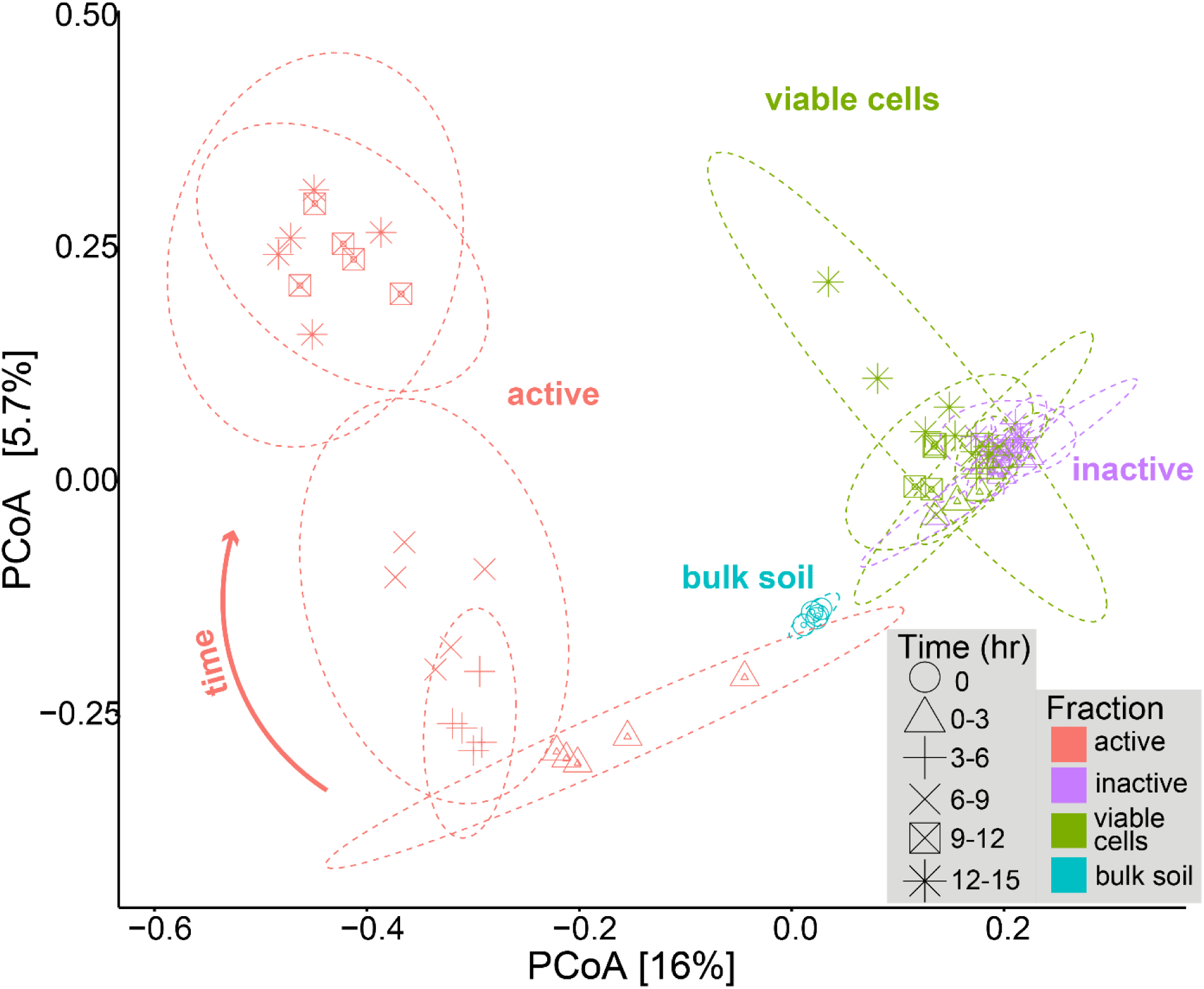
The Bray-Curtis dissimilarity metric PCoA ordination plot showed significant shifts in active community composition over time and a strong divergence between the active and inactive communities. Ellipses represent 95% confidence intervals, colored by fraction and separated by time point. Points represent individual samples, colored by fraction and shaped by time. The community from the bulk soil is time point 0 since this is the soil condition before incubation.

Moreover, the active community shifted over time (symbol type), reflecting a change in community composition, whereas the inactive community showed very little change over time (Figure 3). Interestingly, despite the active community representing less than 1% of all viable cells, the active fraction pulled the entire community toward each timepoint’s respective cluster. A PERMANOVA analysis revealed that in the active fraction, the time point was a significant predictor of the variation in community composition (F = 7.70, *p = 0.001,* df = 1) and explains 25% of the variance. Although the relationship between time and the active community composition is clear in Figure 3 the inactive fraction community composition also had a significant effect of time (F = 1.11, *p = 0.015,* df = 1), but only 5% of the variance was explained by this factor. Even though both fractions had a significant shift in composition over time, the magnitude of the shift was far greater in the active fraction. It’s likely that the shift observed in the inactive fraction is driven by organisms changing their activity status and leaving the inactive fraction.

Overall, 45 phyla were detected, but only 13 were present above 1% relative abundance in the active fraction(Figure 4). Acidobacteriota, Chloroflexi, Proteobacteria, and Plantomycetota were abundant in both fractions. The Firmicutes phylum was highly abundant in the active fraction (Figure 4). The most abundant active ASV was in fact a member of the Firmicutes phylum, classified under the genus *Chungangia*.

**Figure 4:**
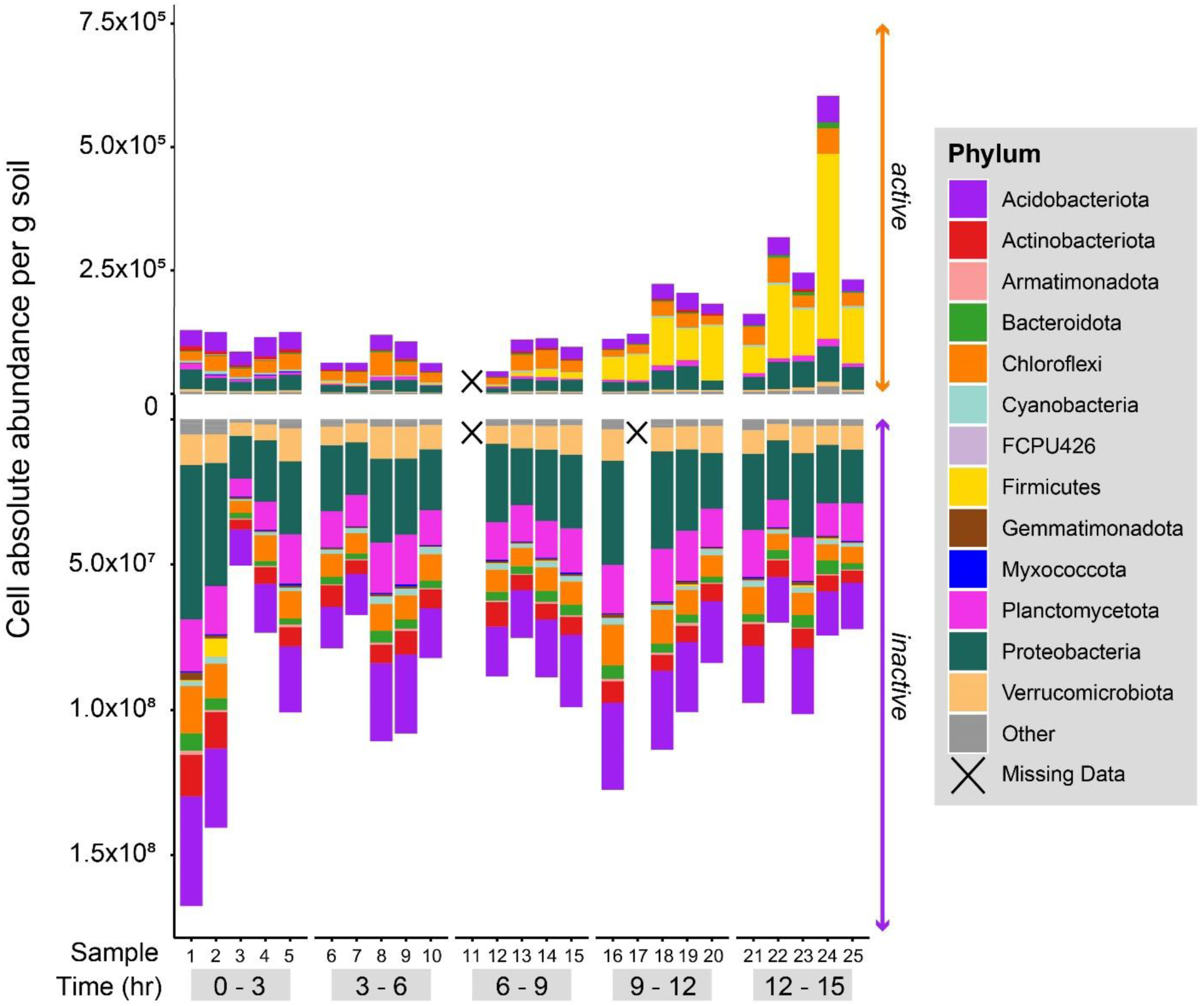
Community composition (absolute abundance) over time at the phylum level and broken down by sample. The top chart shows the active community taxonomic composition, and the bottom shows the inactive community. Axes are on different scales so the active fraction can be seen in detail. Each time point had 5 replicated samples that we destructively harvested, so each sample corresponds to a different microcosm. Selected phylum all have greater than 1% relative abundance in the active fraction (13 out of 45 phyla), other phyla combined abundance is displayed as “Other”.

Using the Boolean Least Squares method included in the mEQO package, we identified a cohort of 8 ASVs whose abundance of active cells per gram of soil correlated best with N_2_O flux rates. This model selected ASVs 1278, 1317, 1342, 1550,1742, 1883, 2556, and 4457 (Table 1, Figure 5). The combined abundance of this ensemble accounted for 95% of the variance in N_2_O flux rates (Figure 5). We also ran this analysis to compare cohorts of different sizes. Particularly, we wanted to see if a cohort of multiple ASVs would correlate better with N_2_O flux rates than a single ASV. The single species model performed worse and only explained 73% of the variance in N_2_O flux rates. The AIC values for the single species model and 8 species model were 42.9 and 1.32, respectively.

**Figure 5:**
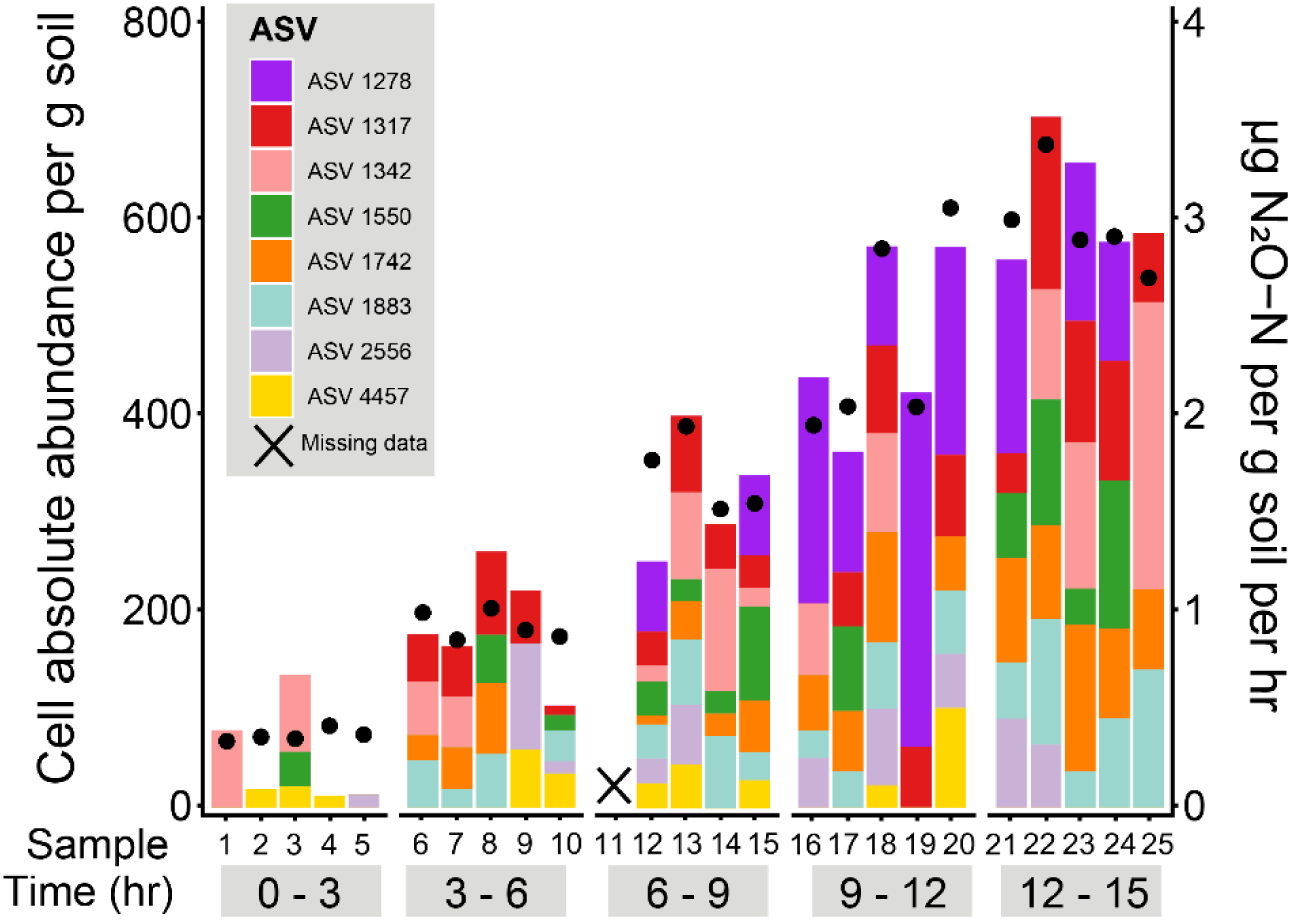
Combined abundance of the active cells of the 8 ASVs identified by mEQO that best correlates with N_2_O flux rates. Each bar shows the combined absolute abundance (left y-axis) of each ASV in the sample. If the ASV is not shown, then it was not present in the sample. Black dots indicate the N_2_O flux rate (right y-axis) measured in each sample. This plot shows that a small number of ASVs (8) combined abundance correlates with N_2_O emission, and that their relative abundance varies among biological replicates, suggesting that there are multiple ASV combinations within this ensemble that explain N_2_O fluxes well.

**Table 1:** Taxonomic classifications from the Silva v138 database for 9 ASVs.

| ASV | Model Type | Phylum | Class | Order | Family | Genus |
| --- | --- | --- | --- | --- | --- | --- |
| ASV109 | Single species | Firmicutes | Bacilli | Bacillales | Planococcaceae | Chungangia |
| ASV1342 | Eight species | Acidobacteriota | Vicinamibacteria | Vicinamibacterales | NA | NA |
| ASV1278 | Eight species | Acidobacteriota | Blastocatellia | Pyrinomonadales | Pyrinomonadaceae | RB41 |
| ASV1317 | Eight species | Acidobacteriota | Blastocatellia | DS-100 | NA | NA |
| ASV1550 | Eight species | Bacteroidota | Bacteroidia | Flavobacteriales | Flavobacteriaceae | Flavobacterium |
| ASV1742 | Eight species | Acidobacteriota | Blastocatellia | DS-100 | NA | NA |
| ASV1883 | Eight species | Acidobacteriota | Blastocatellia | DS-100 | NA | NA |
| ASV2556 | Eight species | Chloroflexi | Anaerolineae | SBR1031 | NA | NA |
| ASV4457 | Eight species | Acidobacteriota | Subgroup 20 | NA | NA | NA |

There was no overlap of ASVs between the 1 species model and 8 species model. To better understand how these groups are different, we used their Silva taxonomy and BLAST results to estimate the functional potential of the organisms identified. Of the 9 ASVs identified between both models, 3 of 9 were classified down to genus, 5 of 9 could only be classified down to order, and 1 of 9 could only be classified to their class. (Table 1)

### 3.4 Microbial community functional potential predicted from taxonomy

Using PICRUSt2 to predict the gene content of the organisms in our study, we found that many nitrogen cycling genes, particularly for denitrification processes, tend to be higher in relative abundance in the active fraction.

We looked for changes in the nitrogen fixation gene, *nifH* between fractions and over time. The predicted relativ abundance of organisms containing the *nifH* gene was significantly lower in the active than the inactive fractions (p < 0.05), and it did not change significantly over time in either fraction. We probed inactive and active fractions for nitrification genes (*amoA, amoB, amoC, hao*). These genes were low in relative abundance in both fractions (less than 0.0001% relative abundance, figure 6), and *amoA* was not predicted to be present in any sample (see methods). None of these genes were significantly different in relative abundance between inactive and active fractions, and neither fraction changed significantly over time (all p-values > 0.05) except *hao* which was significantly different in relative abundance between fractions (p < 0.05).

**Figure 6:**
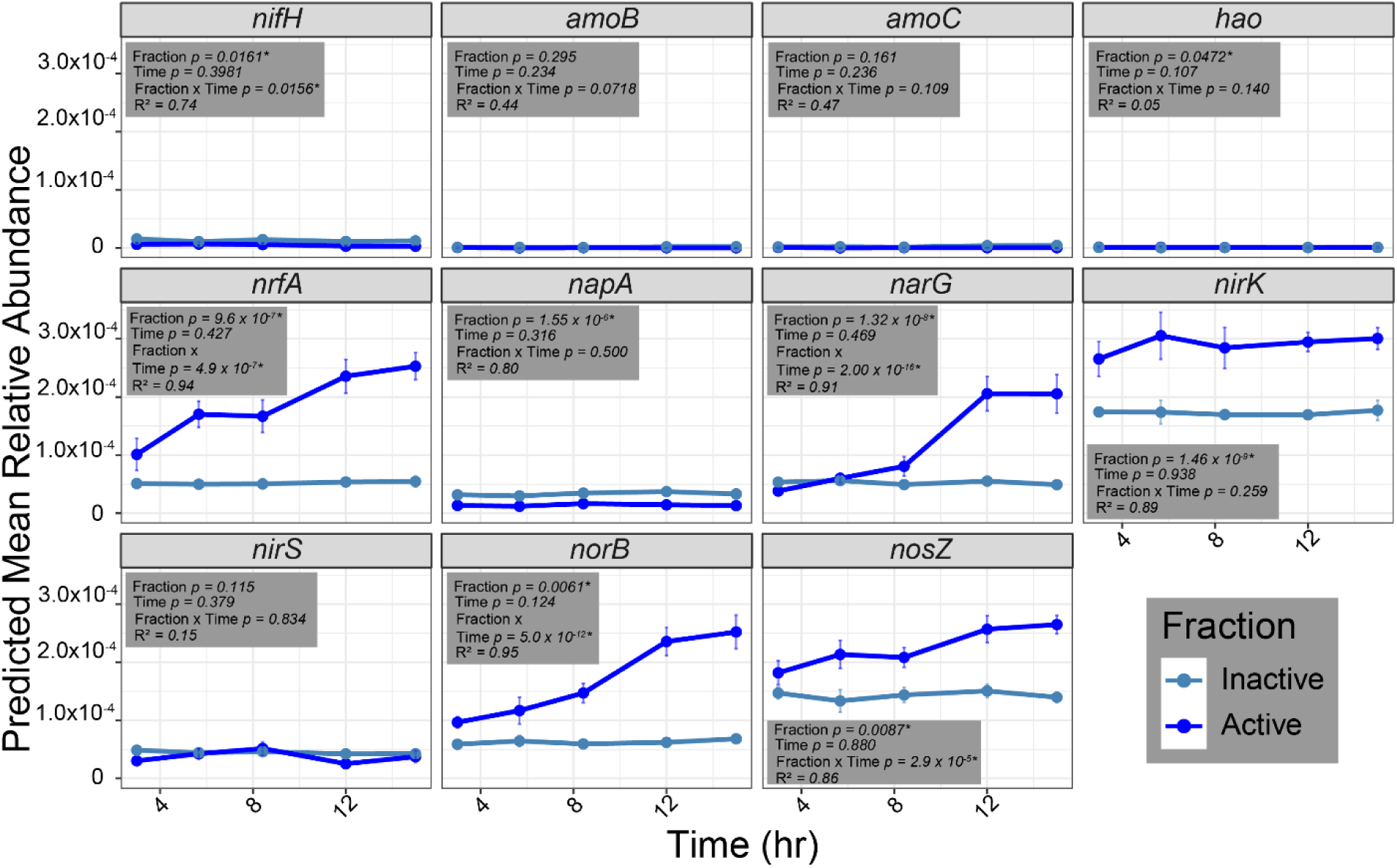
Relative abundance of nitrogen cycling genes predicted by PICRUSt2 across time and separated by activity fraction. Nitrogen cycling genes included are *nifH* (nitrogenase)*, amoB*(ammonia monooxygenase subunit B)*, amoC* (ammonia monooxygenase subunit C)*, hao* (hydroxylamine dehydrogenase)*, nrfA* (nitrate reductase, ammonia forming) *, napA* (nitrate reductase)*, narG* (nitrate reductase)*, nirK* (nitrite reductase)*, nirS* (nitrite reductase)*, norB* (nitric oxide reductase)*, and nosZ* (nitrous oxide reductase). Boxes include p-values where stars indicate significance. Model details are explained in methods section 2.8.

The gene for dissimilatory nitrate reduction to ammonia (DNRA) metabolism, *nrfA*, was significantly higher in relative abundance in the active fraction (p < 0.001) and did significantly increase over time in the active fraction (p < 0.001), but not in the inactive fraction (p = 0.427).

Overall, we found that organisms predicted to contain denitrification genes were more enriched in the active fraction than the inactive. Organisms with nitrate reductase genes, *napA* or *narG* responded differently to incubation conditions. Organisms predicted to contain the *napA* gene were significantly higher in relative abundance in the inactive fraction (p < 0.001) but did not change in relative abundance over time in either fraction (inactive: p = 0.316; active: p = 0.500). However, relative abundance of ASVs predicted to contain *narG* was significantly different across fractions (p < 0.001) and did increase significantly over time (p < 0.001). The relative abundance of organisms predicted to contain *nirK, norB, or nosZ* genes was significantly higher in the active fraction (*nirK:* p < 0.001; *norB*: p < 0.01; *nosZ*: p < 0.01), but only the *norB and nosZ* organisms significantly increased in relative abundance over time (both: p < 0.001). Inactive organisms predicted to contain *nirK, nirS, norB,* and *nosZ* did not change significantly over time (all: p > 0.05).

For the nine ASVs identified by both mEQO models, Table 2 shows the presence and absence of denitrification genes. The single species model ASV, ASV 109, is predicted to have a complete set of denitrification genes with *narG, nirK, norB,* and *nosZ.* Each of the species in the eight species model ASVs are predicted to have one or two denitrification genes. This set of ASVs has nitrate, nitrite, and nitrous oxide reductase genes, but lacks nitric oxide reductase genes.

**Table 2:**
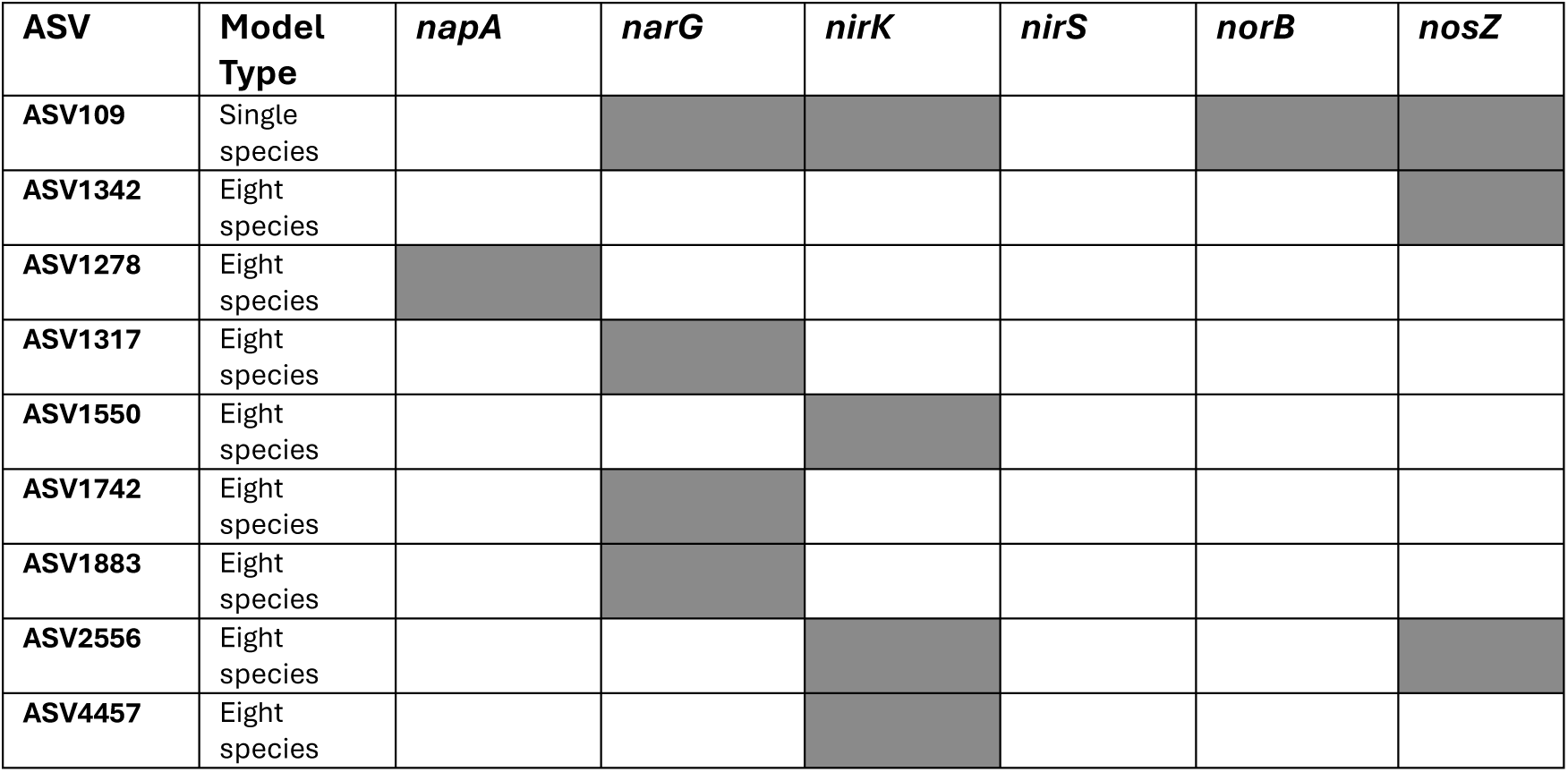
Denitrification gene presence in mEQO ASVs.

## 4 Discussion

Using the BONCAT-FACS method, we are the first to probe translationally active cells during nitrous oxide production in soil. While other studies have applied BONCAT-FACS to soil, biocrust, and activated sludge (Couradeau et al., 2019,Harris et al., 2025, Trexler et al., 2023, Du and Behrens, 2021), and some studies use other methods like transcriptomics to measure denitrifier activity (Brenzinger, Dorsch, and Braker, 2021), our study allows us identity a unique consortia of microbes most associated with N_2_O fluxes. These results show that while N_2_O fluxes increased, the number of active cells and their activity increased, as indicated by their median fluorescence intensity. Additionally, the active microbial community displayed distinct changes in taxonomic composition over time, while the inactive fraction remained relatively stable. The microbial community analysis identified 9 ASVs for further investigation from the mEQO analysis, which allowed us to identify an ensemble of ASVs that correlates best with nitrous oxide flux rates. Using PICRUSt2, we found that organisms containing genes for denitrification tended to increase in relative abundance during the incubation period in the active fraction, and that the ensemble of 8 ASVs identified in the mEQO analysis are predicted to have partial denitrification pathways. Testing this new methodological framework emphasizes that denitrification is a community trait and that a distinct focus on the active community, rather than the entire community, reveals strong and previously obscured relationships between the microbial community and process rates. This work takes a necessary first step in integrating methods, including physiological probing and phenotype-based separation, to directly link microbial functions to emergent soil properties (Hatzenpichler et al., 2020).

### 4.1 The active microbial community during high N_2_O production is taxonomically distinct from the inactive portion

The ordination of the Bray-Curtis dissimilarity metric shows that the composition of the active microbial community shifts through time (Figure 3). The communities change substantially with time early in the incubation, but show less change from 9-12 hours to 12-15 hours of incubation. In this study, we observed a significant and consistent community response to anoxic soil conditions within 15 hours by focusing on changes in only the active community.

Although we see consistency in community taxonomic structure across replicates over time at the phylum level (Figure 4), the mEQO analysis shows some variation between replicates at the ASV level. Despite samples having relatively consistent N_2_O fluxes over a given sampling time (Figure 5), a different consortium of ASVs can result in a similar flux level (Figure 5). These combinatory variations in selected ASVs by mEQO are explained by the fact that the BLS algorithm is selecting an ASV ensemble that correlates best with N_2_O flux rates across samples while a given ASV may not all be in every sample.

In the incubations, the pulse of carbon and nitrogen in conjunction with anaerobic conditions and slight warming (25 °C) can be seen as a disturbance. In the context of disturbance, Martiny and Allison, 2008 define functional redundancy as one taxon with the same process rate as another, and functional similarity as 2 communities with different compositions, but the same process rate. Their analysis found that many microbial communities of different compositions are not functionally similar and functional redundancy between taxa is not common, though other studies have found denitrifier communities to have context dependent functional redundancy (Brenzinger et al.,2015, Salles et al., 2012, Wertz et al., 2006). Assuming the 8 ASVs we identified are contributing to the flux of N_2_O, our results provide a contrasting example, suggesting that functional similarity may be a characteristic of communities generating N_2_O. Indeed, there was predicted to be some functional redundancy across these 8 ASVs (Table 2). The associated predicted functions are distributed across samples, and most functions are associated with the initiation of denitrification. With the high initial nitrate concentrations and the exhausted nitrate supply, the concentration of initiation genes in the identified cohort is not surprising as these are likely denitrification initiators (Pold et al., 2025, Blackmer and Bremner, 1978).

#### 4.1.1 Microbial level of activity correlates better with N_2_O fluxes than percentage of active organisms

Our results show that the level of activity (as indicated by the log of the median fluorescence of each sample, Figure 2B), or the amount of proteins translated, correlated better with nitrous oxide production rates rather than the percentage of active organisms. Since microorganisms can continue translation and cell growth without replication (Munoz-Gutierrez et al., 2024, Gefen et al., 2013), it’s possible that some of the active organisms were translating many proteins without replication. This would then explain why the level of activity correlated better with nitrous oxide production rates. There may be some replication occurring, but our total cell counts over time did not change indicating there was no replication or replication rates matched death rates.

### 4.2 Increasing N_2_O flux rates are correlated with community activity change, rather than changes in the activity of one specific organism

The mEQO analysis identified an ensemble of 8 ASVs that correlates best with the N_2_O flux rate, enabling a test of our second hypothesis; whether 1 ASV or multiple ASVs would correlate with N_2_O fluxes better. Results are consistent with our hypothesis; the increasing N_2_O flux rates in our samples correlated better with the shifting abundance of an ensemble of organisms than with the dramatic growth of one or a few organisms. Work from Pold et al. (2025) corroborates this result as well. In a comparative analysis of denitrification genes in genomes, metagenomes and meta transcriptomes collected from organisms across the globe, they found that partial denitrification pathways are more common than complete pathways suggesting that denitrification is more of a community effort rather than being driven by an individual organism.

None of the identified organisms come from the Proteobacteria phylum, which is known to contain many denitrifiers. (Coyotzi et al., 2016, Kersters et al., 2006, Lycus et al., 2017, Thomsen et al., 2007, Xiang et al., 2023) even when present in the active community (Figure 4). Though incomplete denitrification pathways outnumber complete (Pold et al., 2025), Proteobacteria tend to have complete denitrification pathways (Green et al., 2010, Lycus et al., 2017), which may explain why we did not capture any in our mEQO analysis that potentially identified an ensemble of incomplete denitrifiers.

Using 16S rRNA gene microbial taxonomic analysis, we can only infer the relationship between taxonomic structure and functional potential (Douglas et al., 2020, <u>Aßhauer</u> et al., 2015, Louca et al., 2016). Our analysis shows that the active community is likely to contain a mix of denitrfiers and other anaerobes or facultative anaerobes. We would need to conduct a metagenomic analysis to achieve higher resolution of functions in our community, which is outside the scope of this paper. Regardless, 16S rRNA gene amplicon data can provide preliminary insight into how the active community taxonomic structure may relate to functions. We analyzed 9 ASVs (Table 1) from the active community to predict their functional potential based on taxonomic proximity with cultivated organisms. ASVs were selected by the mEQO tool, both a single species model and eight species model.

#### 4.2.1 Single species model

In the single species model, ASV109, from the phyla Firmicutes, was identified as the one organism whose abundance per gram of soil correlated best with N_2_O flux rates. Interestingly, we found ASV109 was classified differently between the Silva taxonomy database and BLAST. The Silva taxonomy database classified this ASV in the genus *Chungangia.* Currently, this genus has only one representative, *Chungangia koreensis* (Kim et al., 2012). This genus was also the most abundant genus across all samples in the last two time points and represents a large fraction of the Firmicutes bloom seen in these samples (ASV 1 and 2 combined average relative abundance at time point 12-15 is 1.86 ± 0.39%). Kim et al. (2012) describes the representative species as a strict aerobe isolated from marine sediments. Although we did not measure oxygen concentrations throughout the incubation, we expect the replacement of headspace oxygen with nitrogen prevented strictly aerobic metabolic activity. Considering this genus increased in abundance towards the end of the incubation, where oxygen would be even more limited (if present at all), we suggest that this experiment supported the activity of organisms related to the *Chungangia* genus that are metabolically different from the type strain. Two of the top five hits when we BLAST ASV109 showed 99% and 98% similarity to our sequence and were identified to be in the genus *Bacillus.* Based on the literature, *Bacillus* could be a more likely classification for these organisms as members of this genus display a wide variety of metabolic potential. In general, this genus offers a wide range of direct and indirect plant growth promoting metabolisms including nitrogen fixation, phosphorus solubilization, potassium solubilization, and more (Saxena et al, 2019). Additionally, many species have dissimilatory nitrogen metabolisms. Verbaendert et al. (2011) identified 45/87 *Bacillus* species that genes for dissimilatory N reduction and 19 of those were classified as denitrifiers. Indeed, ASV109 was predicted to have a complete set of denitrification genes (Table 2). Further analysis, including genome scale single copy marker genes surveys and effort of cultivation will be needed to fully understand the phylogenomic placement of these ASVs and their metabolic capabilities.

#### 4.2.2 Eight species model

The 8 species model performed better than the single species model when correlating nitrous oxide flux rates with the ASV absolute abundance. ASV 1278 and ASV 1550 were classified by the Silva database to the genus level: RB41 and Flavobacterium, respectively. *RB41* is a genus in the Acidobacteria phylum and has been recognized as responsible for large fractions of carbon flow through a system in aerobic conditions (Stone et al, 2021). Stone et al. (2021) found that RB41 in a small cohort of other organisms (six total) accounted for >50% of carbon cycling. In addition to carbon cycling, *RB41* has been identified as highly abundant in the bulk sludge of and activated sludge bioreactor indicating it may facilitate or play a role in denitrification (Yu et al., 2023). *Flavobacterium* has also been identified from activated sludge, and this genus has a full set of denitrification genes (Conthe et al, 2018), although, ASV1550 was only predicted to contain the *nirK* gene (Table 2). Conthe et al. (2018) indicated that species in this genus tend to be highly competitive N_2_O reducers. Other studies show that *Flavobacterium* abundance tends to be negatively correlated with N_2_O emissions (Ye et al., 2024). We do not see this trend in our samples which may be explained by the differences in experiment length. We incubated samples for up to 15 hours, where Ye et al. (2024) ran their experiment for 77 days.

Five of the eight ASVs were classified to order using the Silva taxonomy. Three ASVs (ASV1317, ASV1742, and ASV1883) were classified as the order DS-100. ASV 1342 and ASV 2256 were classified as Vicinamibacterales and SBR1031, respectively. The order DS-100 has been recovered from wastewater treatment plants (Kristensen et al, 2021). Some of these organisms may be polyphosphate accumulating organisms. The order SBR1031, from class Anaerolinae, likely contains heterotrophic anaerobes (Wang et al., 2018; Trinh et al., 2025) and has been identified in anammox reactors. These conditions suggest these ASVs have both anaerobic and nitrogen-related metabolisms. Another study collected SB1031 organisms from anammox and activated sludge reactors (Bovio-Winkler et al., 2023).

Bovio-Winkler et al. (2023) found that one SB1031 metagenome assembled genome (MAG) had nitrite and nitric oxide reductase genes, though our PICRUSt2 analysis predicted it contains nitrite and nitrous oxide reductase genes.. Additionally, all their recovered MAGs from SB1031 could break down complex polymers through fermentation. Finally, the order Vicinamaibacterales has been associated with phosphorus solubilization and also positively correlated with nitrate concentrations (Wu et al., 2021).

The last organism identified in our mEQO analysis, ASV 4457, could only be classified down to its class, subgroup 20 from Acidobacteria. Thereis little literature to describe subgroup 20. Generally, the Acidobacteria phylum is considered very diverse, hosting many different functions and metabolisms including nitrogen metabolisms (Kalam et al., 2020). Other studies have pointed out that organisms in the Acidobacteria phylum may play a central role in N-cycling, with denitrification capabilities, and C-cycling (Ward et al., 2009; Banerjee et al., 2016; Banerjee et al 2018).

Based on the functional potential these eight ASVs have, we see that the group of organisms that correlate best with N_2_O flux rates may be taking part in a broader suite of metabolisms than just those related to denitrification. While we expected and found that several of the organisms come from groups known for denitrification, this analysis also highlights the role of other nutrient cycling in supporting N_2_O fluxes, which can help explain the spatial and temporal variation in these flux rates seen in field conditions. Here, we suggest that denitrification is a community trait supported by a suite of organisms performing additional metabolisms. This is evident when we compare the maximum potential rate of N_2_O production based on NO_3_^-^-N consumption rates (5.13 µg N_2_O-N g dry soil^-1^ hr^-1^) to the actual peak N_2_O production rate (3.84 µg N_2_O-N g dry soil^-1^ hr^-1^). This deficit of 1.29 µg N_2_O-N g dry soil^-1^ hr^-1^ that could be produced indicates that some nitrogen is being used in other processes, NO_3_^-^-N is being immobilized, or N_2_O-N is being reduced to N_2_. ThePICRUSt2 analysis predicts a relative abundance of organisms with *nosZ* that is consistent with the relative abundance of other organisms containing N-cycling genes suggesting denitrification was possibly a complete pathway. N_2_O priming hypotheses would not explain this deficit as the four current hypotheses rely on N limitation or exhaustion of the input, which was not reached (figure 1) or do not explain a negative effect on N_2_O emissions, but rather a decrease in soil organic matter derived N_2_O emissions (Daly et al., 2024, Blagodatskaya and Kuzyakov, 2008). Dissimilatory nitrate reduction to ammonia (DNRA) cannot fully explain the deficit as the maximum nitrate utilization rate by organisms with DNRA metabolisms is unlikely to exceed 0.16 µg NO_3_^-^-N g dry soil^-1^ hr^-1^ based on the rate of ammonium accumulation. Previous studies have inconsistently correlated taxonomy to rates of N_2_O (Bahram et al., 2022, Butterbach-Bahl et al.,2013, Lee and Kang, 2015, Salles et al., 2012), but by isolating the active fraction, we see a consistent relationship between the microbial community, predicted relative abundance of organisms with denitrification genes, and N_2_O emissions (figure 5, figure 6).The ASV from the single species model was not included in the 8 ASV model. Although the 8 ASV model is an ensemble of organisms whose abundance correlated best with the N_2_O production rate, suggesting they were facilitating or participating in denitrification, they were likely not the only organisms in the microcosms denitrifying. Broadly, the changes in predicted denitrification gene relative abundance during the incubation indicate that a larger fraction of the community was engaged in denitrification. Specifically, members of the Firmicutes phylum have indeed been linked to denitrification and dissimilatory reduction of nitrate metabolisms(Anderson et al., 2018, Lycus et al., 2017). The work from Pold et al. (2025) suggests that these Firmicutes could be initiators (defined as organisms with just *nir,* nitrite reduction, genes or *nir* and *nor,* nitric oxide reduction, genes) of denitrification or complete denitrifiers due to their rapid growth in our experiment. Based on ASV109’s, a Firmicutes, predicted complete set of denitrification genes (Table 2), it’s likely that they were conducting complete denitrification. Being a complete denitrifier could explain why no Firmicutes species, or the ASV we identified in our single species model, are in our 8 species model since these organisms are not reliant on others organisms for substrates so their abundances are less linked to other organisms.

#### 4.2.3 Firmicutes as a dominant phylum across later time points

In all of the samples during the last 2 time periods, 9-12 hours and 12-15 hours, the Firmicutes phylum dominates the active microbial community as the most abundant phylum (43.0% ± 10.5; 45.2% ± 11.3) and drove a significant difference in Pielou’s evenness (Figure 4, Supplementary Figure 3). Interestingly, none of the 8 organisms identified by the mEQO analysis are from the Firmicutes phylum, and none of the most abundant ASVs were part of this cohort either. ASV109 is from the Firmicutes phylum and was the best single ASV for explaining N_2_O flux rates (discussed in section 4.2.1).

One study, (Qi et al., 2021) found that abundance of the Firmicutes phylum increased following glucose C additions and with narrower soil C:N ratios. Since our soil amendment had a C:N ratio of 14:1, which is not particularly low compared to a typical agricultural soil C:N (our soil C:N is ∼10), but lower than many plant residues added to soil, this could stimulate an increase in Firmicutes abundance. Another study on biocrusts reports a similar bloom of Firmicutes not in response to carbon, but rather rehydration. Karaoz et al., 2018 found that 18 hours after rehydration, Firmicutes, particularly in the *Bacillales* order, increased significantly. We also found that Firmicutes from the order *Bacillales* bloomed within a similar timeframe.

Firmicutes is a diverse phylum (Galperin et al, 2013) that includes spore formers, non sporeformers (which likely lost the ability to form spores as an adaptation to nutrient rich conditions), aerobes, and anaerobes. Given the nearly undetectable levels of Firmicutes in the first two time periods of our incubation (0 - 3 hr, 3 - 6 hr), it’s likely that the Firmicutes we observed were r-strategist spore forming anaerobes that take advantage of high nutrient, anaerobic conditions. Since this is such a diverse phylum, we cannot infer the exact functions associated with the organisms that bloomed, but it is likely that they are complete denitrifiers (Table 2) Additionally, this same phenomenon has been observed in another study where high Firmicutes abundance was linked to high denitrification enzyme activity (Anderson et al., 2018)

To further explore the active microbial community and their functions, metagenome data will need to be collected. Then, we can explore the functions specifically associated, rather can predicted based on 16S amplicon sequences, with the active microbial community and see how they change through time.

## 5 Conclusion

In this study, we aimed to uncover the link between the active microbial community structure and N_2_O emissions from soil. We found that less than 1% of the microbial community was active during N_2_O production and that the percentage of organisms that were active did not correlate as well with N_2_O flux rates as the log of the median fluorescence – a metric associated with protein synthesis. In terms of the microbial community, we found that the active microbial community changed significantly over time, while the inactive microbial community remained relatively stable. Our analysis using the mEQO tool identified an ensemble of 8 ASVs in the active community, of which none were the most abundant in the community, whose combined absolute abundances correlated better with N_2_O flux rates than any single ASVs. Using the taxonomy of these 8 ASVs, we inferred that they were a functionally diverse group, including potential denitrification as well as C-cycling, P-solubilization, and fermentation, which could be facilitating the high rates of N_2_O emissions. This study emphasizes the need for focusing directly on the active microbial community. Since less than 1% of the community was changing their activity status, any changes in their abundance would likely be missed in studies looking at the whole microbial community.

Overall, our results reveal that N₂O fluxes are potentially driven not by changes in a single taxon but by shifting ensembles of active microorganisms whose combined functional potential supports consistent emissions, underscoring the importance of activity-based approaches for predicting microbial contributions to greenhouse gas production. We recognize that these results are highly context dependent as we only tested one soil type. However, this study has proved the utility of this methodological framework that integrates microbial community-based analysis with process rates which warrants further testing across more diverse soil types. This approach has significantly reduced the complexity of the microbial community associated with N_2_O production by constraining our focus to less than 1% of the microbial community. In addition, the mEQO analysis yielded an even smaller subset of the active microbial community that can be specifically analyzed in future studies to discern the mechanistic relationship between these ASVs and N_2_O emissions. The PICRSUt2 analysis revealed that the 8 ASVs and the active community as a whole were predicted to contain denitrification genes and shifts in the active community mimicked shifts in the relative abundance of denitrification genes. This finding supports that denitrification is less of a trait of an individual organisms, but rather a community and will impact how we predict and manage N_2_O emissions.

## CRediT authorship contribution statement

**Jonah Gray:** Conceptualization, Data curation, formal analysis, methodology, visualization, writing – original draft, writing – review and editing. **Jennifer Harris:** Methodology, writing – review and editing**. Jason Kaye**: Conceptualization, methodology, writing – review and editing, funding acquisition, supervision. **Estelle Couradeau**: Conceptualization, methodology, visualization, writing – review and editing, funding acquisition, supervision

## Declaration of competing interests

The authors declare that they have no relationships that could be considered conflicts of interest.

## Acknowledgements

This work was made possible by the Pennsylvania State University Core Huck Institute of the Life Sciences Genomics, Metabolomics and Flow Cytometry Core Facilities and the Pennsylvania State University Agricultural Analytical Services Laboratory. Special thanks to members of the Flow Core Facility, Dr. Rajeswaran Mani and Mitchell Koptchak for guidance during cell sorting and BONCAT-FACS data analysis. Additionally, Brosi Bradely for guidance in soil acquisition and analyses.

## Funding statement

This work was supported by the Natural Resources Conservation Services [grant number: NR233A750004G055, 2023] and by the United States Department of Agriculture hatch appropriations [grant number: PEN04949; grant number: PEN04977].

## Appendix A. Supplementary information Supplementary Methods

## S1.1 Experimental Design

**Supplementary figure 1:**
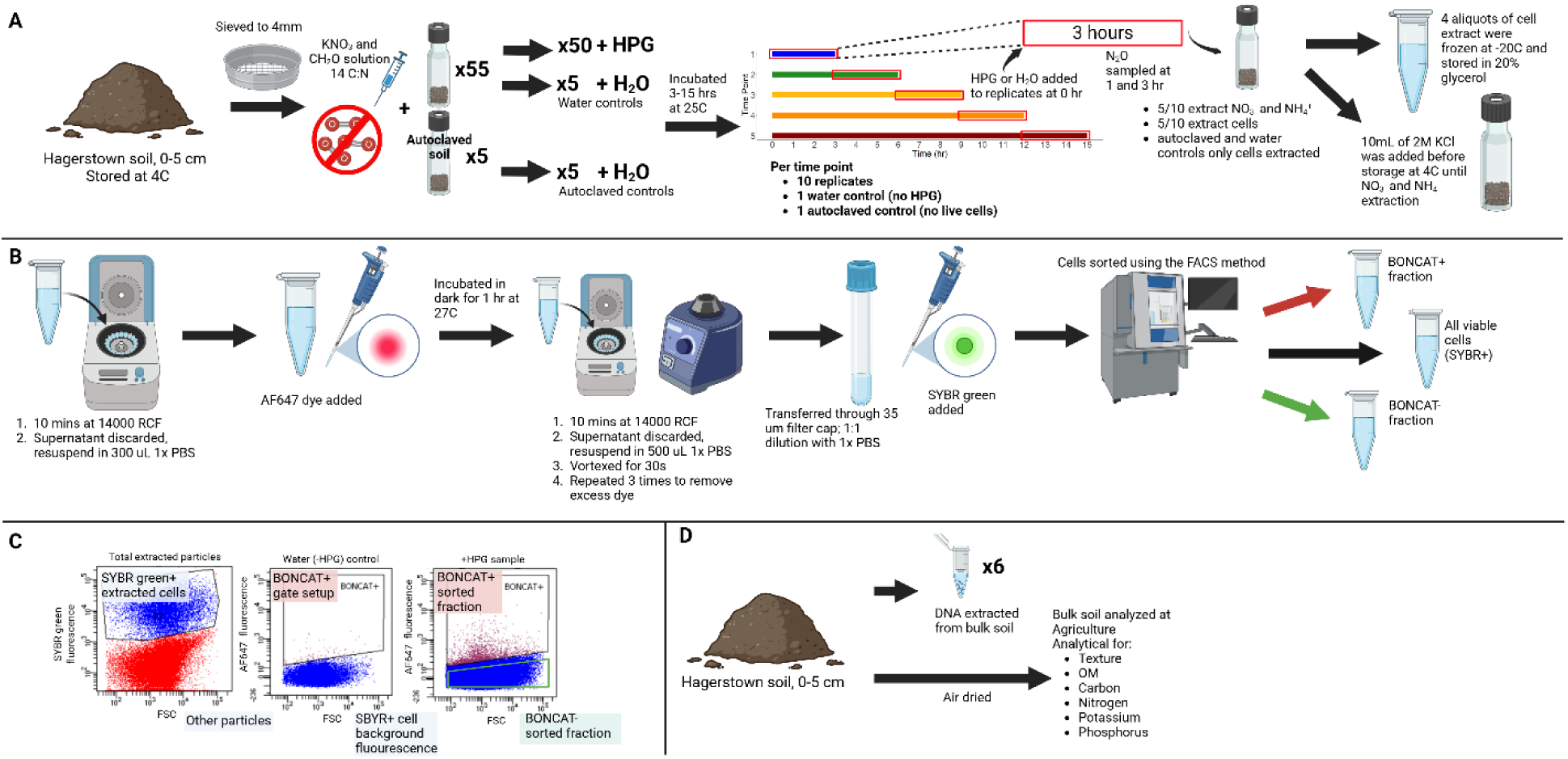
(A) For the incubation, soil was sieved and distributed into exetainer vials. Autoclaved control soil was autoclaved before being added to the vials. Vials were organized into 5 groups and incubated for various lengths of time that increased in 3 hour increments up to15 hours. During the last 3 hours of each group’s incubation (red box), HPG or water was added to the vials and N_2_O was sampled twice. Cells, nitrate and ammonium were extracted from the soil following the incubation. (B) Aliquots of cells extracts were centrifuged at 14000 RCF to pellet the cells and resuspended in 300 µL 1X PBS. A solution containing the AF647 dye was prepared (methods) and added to the vials. They were incubated in the dark for 1 hour. To remove excess dye, cells went through a washing procedure in 1X PBS 3 times. After being transferred to a new text tube, the SYBR green dye was added and samples were diluted. Samples were processed this way in batches of 3 – 7 and each batch was sorted into the 3 fractions (BONCAT +, BONCAT -, all viable cells) on the same day. (C) Cells were sorted using a nested gating strategy. The SYBR gate was determined by autoclaved soil controls with and without the two dyes. Since no cells were present in the autoclaved soil, this allowed us to see the background fluorescence of the soil and other particles. Cells in the SYBR green+ extracted cells gate were passed to the next step to determine activity. Water controls with and without the two dyes were used to draw the BONCAT + gate (active cells). The second panel shows a water control with both dyes. We limited the BONCAT + gate false positive rate to 0.05%. Cells were sorted into 3 fractions BONCAT + (active), BONCAT - (inactive), and SYBR green + (all viable cells). (D) The bulk soil had DNA extracted and was dried and analyzed at the Pennsylvania State University Agricultural Analytical Services Laboratory. Created in BioRender. Gray, J. (2026) https://BioRender.com/whc59lu

## S1.2 HPG solution preparation

To prepare an HPG solution for the microcosm experiment, a stock of 20mM HPG was diluted in Milli-Q water to 4.84 mM. When added to the soil microcosms this concentration results in 1 µmol HPG per gram of dry soil. We opted to use a higher concentration HPG solution to maintain 1 µmol HPG per gram of dry soil without flooding the soil. The HPG solution was filter sterilized into 5 different autoclaved Wheaton bottles the day prior to use. Each bottle was immediately purged of oxygen using N_2_ gas. To ensure dissolved oxygen was purged as well, each vial was purged for 30 seconds, then shaken for 30 seconds. This process was repeated 4 times. HPG was stored in the fridge overnight until the bottles were removed the next morning. Bottles were removed 30 minutes prior to use to allow them to reach room temperature.

Following the experiment, six subsamples were taken from the original soil for drying and freeze drying to determine the methionine content in the soil. Methionine from soils was extracted by adding 5 ml of LC-MS grade water to 1g of soil. Samples were incubated for 1 hour at 4 °C, vortexed for 5 seconds every 15 minutes. Samples were centrifuged at 3220 RCF for 15 minutes at 4 °C. We filtered supernatant with a 0.45 µm filter and freeze dried using a HarvestRight freezer drier. After freeze drying, samples were sent to the Huck Life Sciences Metabolomics Core Facility to test methionine levels in soil following the same method detailed in the supplementary materials of Harris et al., 2025. On average, our soil contained less than 1 nmol/g of dry soil.

## S1.3 Nitrate and Ammonium analysis

Initial nitrate and ammonium extractions were conducted shortly after the soil was collected. Five subsamples were used to calculate average concentrations. Extractions were conducted by combining ∼20 g of soil into a specimen cup with 100 mL of 2 M KCl. They were shaken for 1 hour then removed from the shaker and placed upright to allow sediment to settle for ∼1 hour. Extracts were filtered through Whatman #1 filter paper and collected in scintillation vials for analysis. Nitrate and ammonium concentrations were determined using a spectrophotometer and the Vanadium (III) Chloride method (Doane and Horwath, 2003) and the modified Indophenol Blue Method using salicylate (Utomo et al., 2023).

Five vials from the microcosm experiment had 10 ml of 2 M KCl added immediately after being removed from the incubator. They were stored at 4 °C until they were extracted and analyzed using the procedure outlined above.

## S1.4 BONCAT-FACS: Reaction mixture and cell sorting

To prepare the reaction mixture for cell staining, we prepared it daily based on the number of samples being processed. Per sample, it consisted of 1.125 µl copper sulfate (CuSO4 100 µM final concentration), 2.25 µl tris-hydroxypropyltriazolylmethylamine (THPTA, 500 µM final concentration), and of 0.747 µl AF647-picolyl azide (5 µM final concentration). Reagents were added in order, mixed gently and incubated in the dark for 3 mins. This solution was buffered in 11.25 µl of 5 mM sodium ascorbate freshly prepared in 1X PBS, 11.25 µl of 5 mM aminoguanidine HCl freshly prepared in 1X PBS and 184.5 µl of 1X PBS. It was gently mixed and stored in the dark until use.

To prepare the samples for sorting, each sample, water control and autoclaved control received the reaction mixture. They were placed in an incubator in the dark at 27 °C and shaken slowly at 50 RPM (< 1 RCF) for 1 hour. After the incubation, cells were pelleted at 14000 RCF for 10 minutes. The supernatant was removed and cells were resuspended in 1 mL of 1X PBS. Pelleting and resuspension was repeated 3 times to wash the cells of any excess dye. After the final wash, cells were resuspended in 500 µL 1X PBS and vortexed. Samples were filtered through a 35 μm filter (BD-falcon 5 ml tube with cell strainer cap, Corning Inc., Corning, NY, USA) to remove any remaining large particles that could clog the flow cytometer. Finally, we added SYBR green (excitation: 498 nm; emission: 522 nm) counterstain at a final concentration of 0.5 µM.

During cell sorting, each biological replicate underwent an initial sort that collected vials of active and inactive populations and all viable cells. The cells were sorted into 100 µL of 0.2µm filter sterilized 50% glycerol and immediately frozen and later transferred to long-term storage in a -80 °C freezer. Setting the flow rate to 32 µL/min for 2 minutes on the cell sorter allowed us to obtain approximate cell counts to estimate the amount of active, inactive, and total viable cells per gram of dry soil.

All biological replicates exhibited below 1 percent activity, which resulted in suboptimal conditions for cell sorting. Subsequent sorts were conducted to collect technical replicates and increase the total number of active cells. Additionally, due to limited quantity of cell extract aliquots, we did not have enough sample to reach an ideal minimum threshold 5.0 x 10^4^ cells for all sorted populations for each biological replicate (Reichart et al., 2020). Cell yield ranged from 8.2 x 10^3^ up to 1.2 x 10^5^ cells.

## S1.5 PCR

The MicroGEM prep kit chemically lysed cells to extract their DNA. Sorted cells were pelleted at 20,000 RCF for 10 minutes to ensure most cells would not be in the supernatant. The supernatant was discarded and 5 µL of the MicroGEM extraction mixture was added to cell pellets. Samples were immediately vortexed for 2 minutes and then spun down, then vortexed for 1 minute and spun down two more times. For the chemical lysis to occur, the samples were placed in the thermocycler for 30 minutes at 37 °C and then 30 minutes at 75 °C. Following this process, samples were immediately removed from the thermocycler to be prepared for amplification.

We prepared our PCR mixture using the Invtirogen Platinum Master Mix, a GC enhancer, sterile water, and primers 515F and 806R (Apprill, et al., 2016, Caporaso et al., 2011, Caporaso et al., 2012, Parada et al., 2016). We added 20 µL of the mixture directly to the samples post the MicroGEM procedure. Samples were then placed in the thermocycler for amplification following the program in supplementary table 1. Following amplification, products were frozen at –20 °C until they were sequenced.

**Supplementary Table 1:**
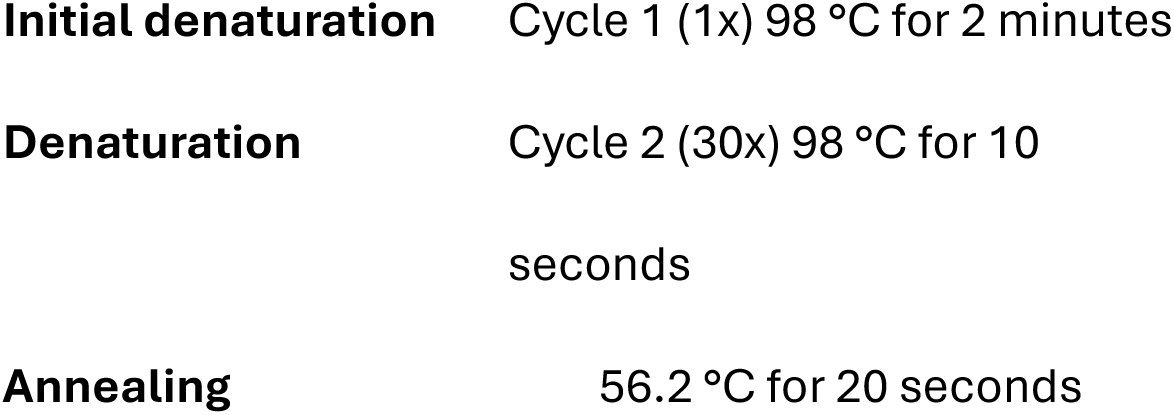
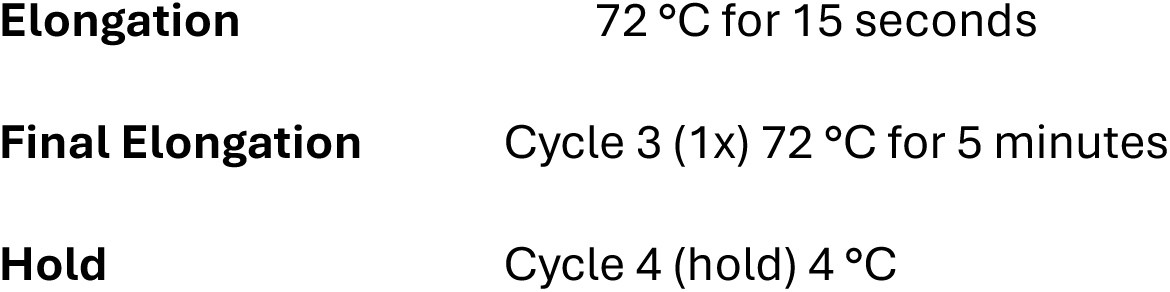
Thermocycler program used for PCR.

## S1.6 16S rRNA gene amplicon data analysis

Data analysis was conducted in R using a DADA2 pipeline (Callahan et al., 2016). 16S rRNA gene sequences were filtered and trimmed, forwards and reverse reads were merged, and chimeras were removed. Before taxonomy was assigned, samples were rarefied to the lowest sequence depth across samples of ∼19K paired reads per sample. We assigned taxonomy using the Silva version 138 database (Callahan et al., 2016, McLaren, 2020, Quast et al., 2013, Yilmaz et al., 2014) and the QIIME 2 2020.6-2021.2 Naive Bayes Classifier for Silva version138 with 99% OTUs full-length sequences (Bokulich et al., 2018, Robeson et al., 2020). The R code is available on GitHub and linked in the data availability statement along with a data frame containing the number of reads per sample through each step of the process.

To estimate the abundance of cells per gram of dry soil, we utilized cell count estimates collected on the Bigfoot Spectral Cell Sorter. Each sample was initially sorted at a constant flow rate of 32 µL/min for 2 minutes. Using the number of cells sorted per volume of liquid, we calculated the cells per µL and used that to find the cell count per gram of dry soil for each sample. To estimate the absolute abundance of the phyla in our samples, we multiplied the phylum’s relative abundance by the sample’s cell count per gram of dry soil. Since the active cell biomass was low, and we had limited replicates, estimates of abundance are therefore based on one aliquot per sample. The rest of the aliquots were needed to maximize our collection of active cells to ensure we had a fair representation of the active community.

## Supplementary results and discussion

### S2.1

**Supplementary figure 2:**
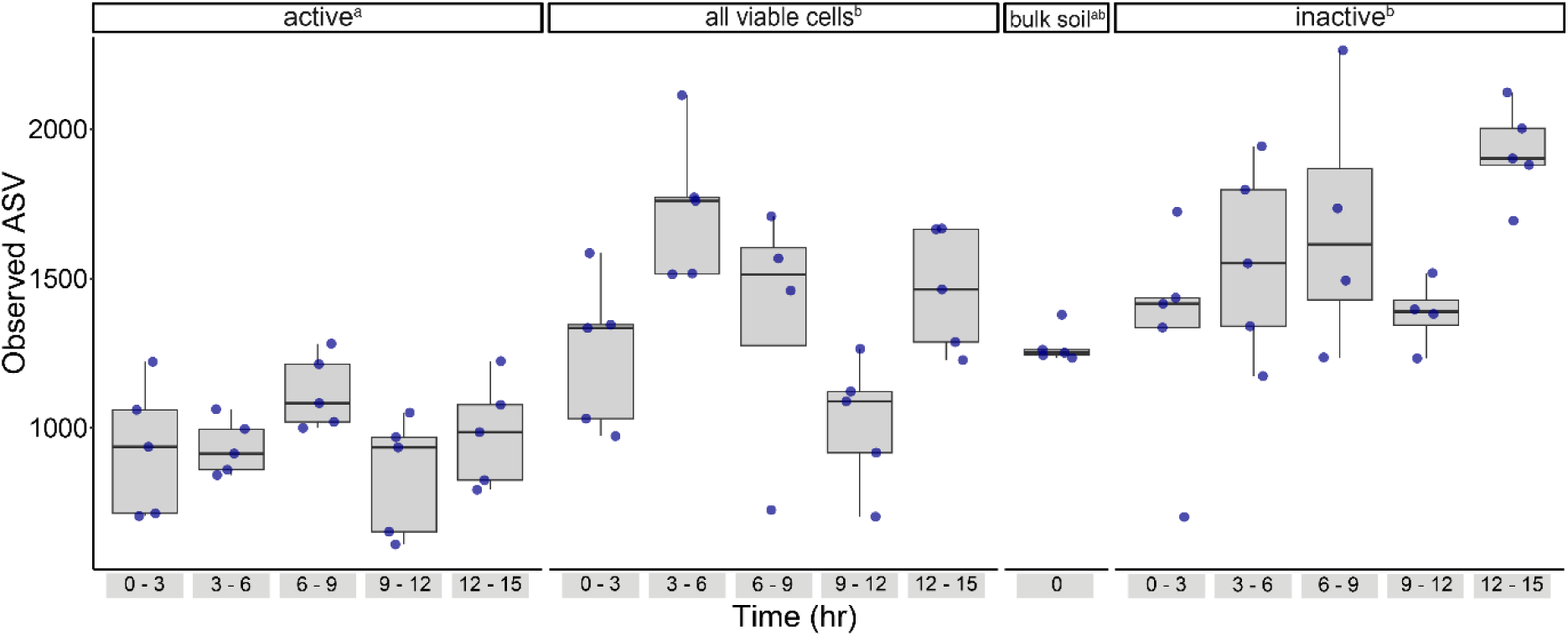
Alpha diversity of the different fractions at different time points. The observed ASVs ranged from 610 to 2265. A one-way ANOVA detected a significant difference between the different fractions (*p < 0.001*), and a Tukey post-hoc comparison showed the observed ASVs in the active fraction are significantly different than from all viable cells and from the inactive fraction (*p < 0.001, p < 0.001*), but none were significantly different than the number of ASVs in the bulk soil. Significant differences among cell types are denoted by superscripted lowercase letters adjacent to cell type labels.

**Supplementary figure 3:**
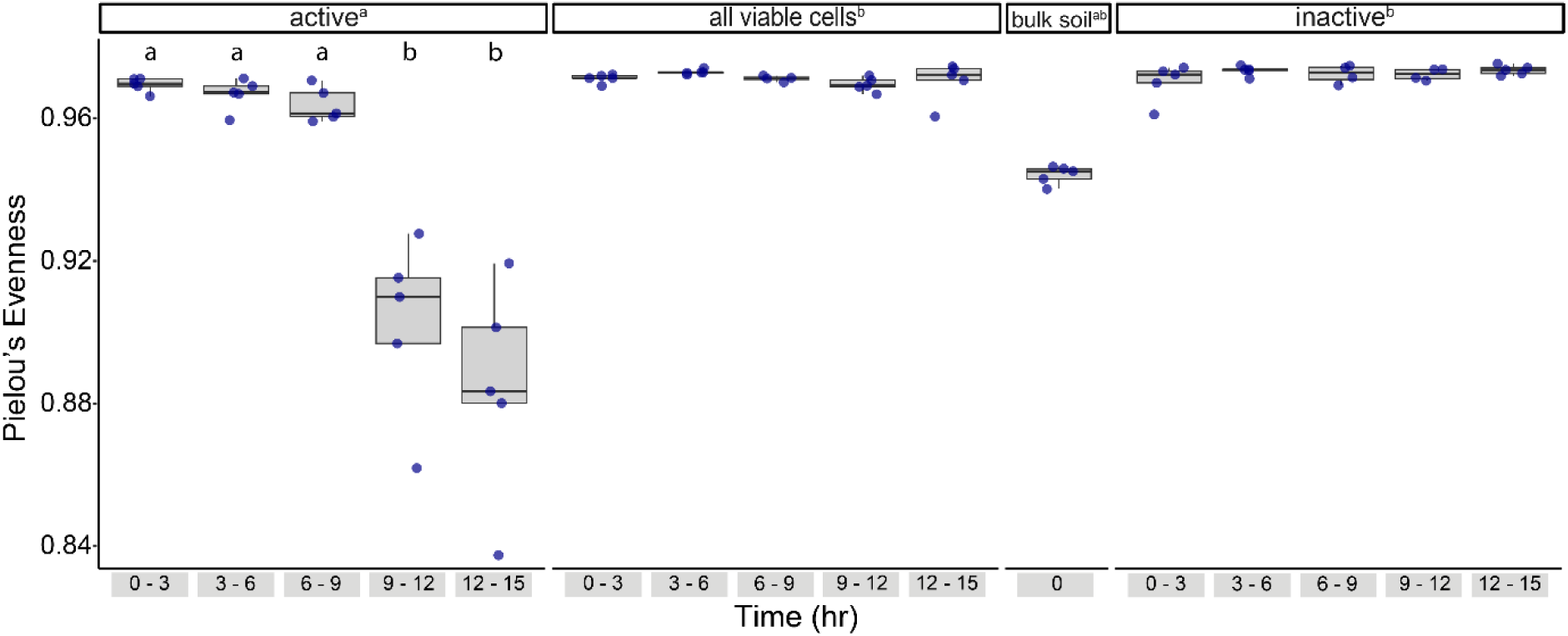
Pielou’s evenness of the different fractions at different time points. A one-way ANOVA detected a significant difference between the different fractions (*p < 0.001*), and a Tukey post-hoc comparison showed the evenness in the active fraction was significantly different from all viable cells and the inactive fraction (*p < 0.001, p < 0.001*), but the bulk soil was not significantly different in evenness than any other fraction. In the active fraction, an ANOVA test showed evenness was significantly different over time (*p < 0.001*), and a Tukey post-hoc test revealed that the last two time points were significantly different than the first three (*4-1, p < 0.001; 4-2, p < 0.001; 4-3, p < 0.001; 5-1, p < 0.001; 5-2, p < 0.001; 5-3, p < 0.001*). The first three time points are not significantly different from each other, and the last two time points are not significantly different from each other. Significant differences among cell types are denoted by superscripted lowercase letters adjacent to cell type labels.

**Supplementary figure 4:**
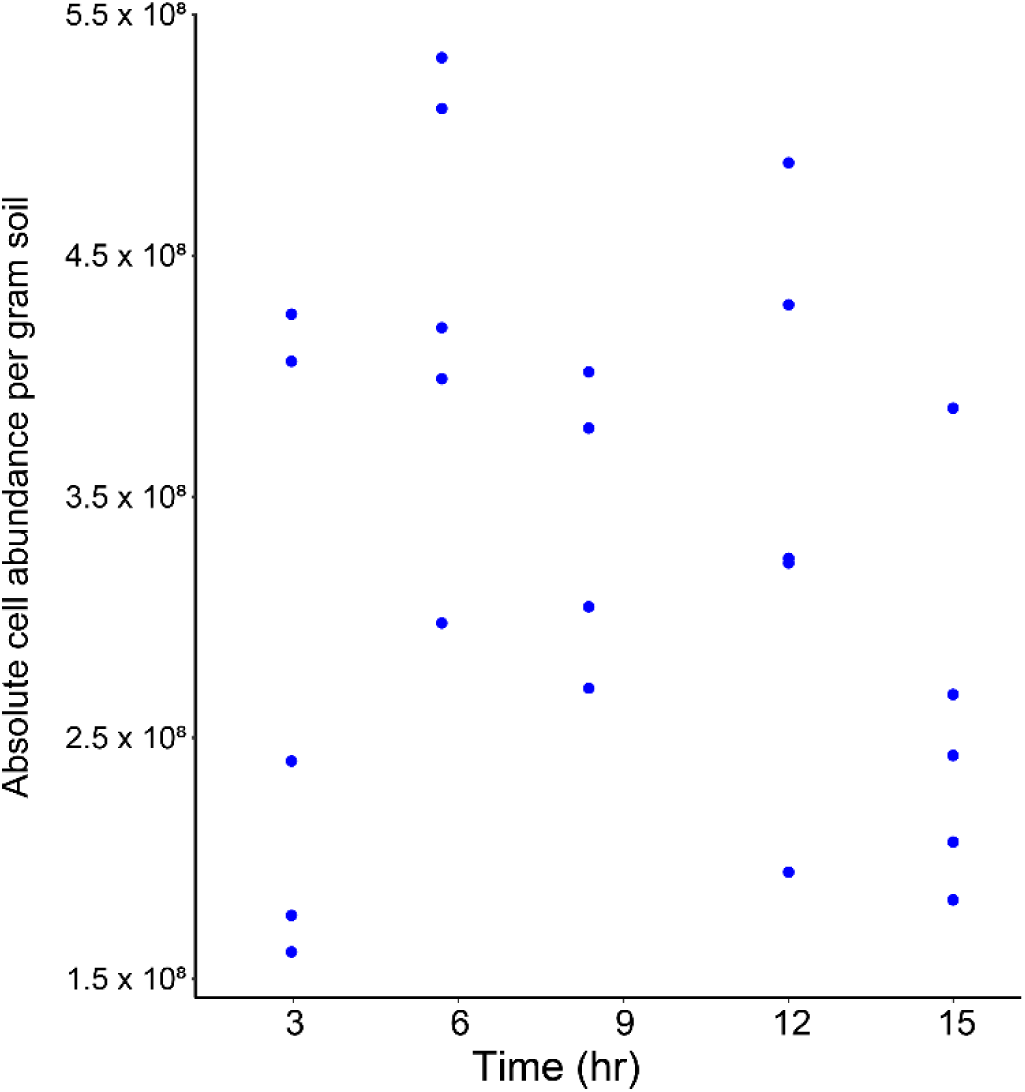
Total cell abundance over time. Cell abundance did not significantly vary with time *(Time vs total cell count, R^2^ = 0.03, p = 0.38)*.

### S2.2 Incubation methods evaluation

#### S2.2.1 N_2_O production rates

The highest nitrous oxide flux rate we collected during hours 13-15 of the incubation - ∼72 mg N/kg soil/day - likely represents a maximum emission scenario. Since we supplied abundant carbon and nitrogen as well as anaerobic conditions, this is not a surprising outcome. In a similar incubation approach, Swerts et al., 1996, supplied the same concentrations of carbon and nitrogen to their soils. They also recorded an N_2_O-N flux rate of ∼72 mg N/kg soil/day. In contrast, Lussich et al., 2024, conducted soil incubations with added cover crops and only reported 2.5-3.0 mg N/kg soil/day. This is far lower than our peak flux rate. Since Lussich et al., 2024 did not supply anaerobic conditions and incubated at a lower temperature (21 °C in their study *vs.* 25 °C in our study), the lower flux rate they observed would be expected. Thus, our study focused on a theoretical maximum rate of N_2_O fluxes by supplying accessible C and N. To better understand how this process could be different in field conditions, we would need to work with more realistic conditions including natural amendments such as plant residues.

#### S2.2.2 NO_3_^-^, NH_4_^+^ and N_2_ transformations

Since we used N_2_ to achieve anaerobic conditions in our microcosms, we expected that there may be some biological nitrogen fixation occurring. Increasing NH_4_^+^ concentrations in the soil are consistent with this possibility. Although, given the magnitude of difference between nitrate used versus the ammonia added, we can conclude that biological nitrogen fixation was not a dominant process and likely only represents a small fraction of the active microbial community. Since nitrogen fixation is a very oxygen sensitive process (Myrold, 2021), the increase in ammonium concentrations suggests that we successfully achieved anaerobic conditions in the microcosms.

Nitrate concentrations decreased consistently throughout the incubation while ammonium increased. Although we cannot be sure no nitrification was occurring, we would expect consistent decreased concentrations of ammonium if this was a dominant process. Additionally, we might see an increase or no change in nitrate concentrations. We did not observe either of these in the microcosms, and nitrification is an aerobic process, so we can conclude nitrification was likely not occurring in our microcosms.

## Data Availability

Analyses were conducted using the R Statistical language (version 4.2.3; R Core Team, 2023). Packages used in data analysis include report (Makowski et al., 2023), ggplot2 (Wickham, 2016), dada2 (Callahan et al., 2016), ShortRead (Morgan et al., 2009), Biostrings (Pagès et al., 2022), phyloseq (McMurdie and Holmes, 2013), corncob (Martin et al., 2025), DESeq2 (Love et al., 2014), microbiome (Lahti and Shetty, 2012-2019), DECIPHER (Wright, 2016), phangorn (Schliep, 2011), tibble (Müller and Wickham, 2023), lme4 (Bates et al., 2015), vegan (Oksanen et al., 2022), emmeans (Lenth, 2025), tidyverse (Wickham et al., 2019), and mEQO (Shan et al., 2023). Raw and processed data is available here, doi: 10.17632/ttvkr8ng3t. Code for reproducibility is available at https://github.com/jonah-n-gray/Active-Microorganisms-during-N2O-production. The sequencing data are available at National Center for Biotechnology Information (NCBI) Sequence Read Archive (SRA), under BioProject PRJNA1358296.

## Notes

### Competing Interest Statement

The authors have declared no competing interest.

